# Repressed mTORC1 signaling and transient dendritic pruning support axonal regeneration

**DOI:** 10.64898/2026.02.11.705324

**Authors:** Anyi Zhang, Luca Masin, Steven Bergmans, Elena Putti, Shahad Albadri, Fabienne E. Poulain, Filippo Del Bene, Lieve Moons

## Abstract

Dendrite degeneration is an early pathological feature following axonal damage, yet its role during successful axonal regeneration remains poorly understood. Using sparse labeling of retinal ganglion cells in adult zebrafish, we show that axonal regeneration is accompanied by a transient phase of dendritic pruning following optic nerve crush. Dendritic pruning occurs during axon elongation and is reversed after brain reinnervation. Although insulin–mTOR signaling is rapidly upregulated immediately after injury, it is subsequently suppressed, coinciding with dendritic pruning and sustained axonal growth. Prolonging mTOR activation through insulin treatment or direct pathway stimulation reduces dendritic pruning in an mTORC1-dependent manner but severely delays axonal regrowth. Together, these findings identify dendritic remodeling as a key temporally regulated component of successful axonal regeneration and reveal a functional trade-off between dendritic preservation and axon regrowth. This balance may represent a critical constraint for promoting CNS repair in mammals.

## Introduction

The adult mammalian central nervous system (CNS) has a limited capacity for repair. Axonal damage, resulting from trauma or chronic degeneration, leads to loss of function and eventual neuronal death^1–3^. Dendritic degeneration is a well-established early response to axonal damage in CNS neurons and is thought to contribute to further decline of neuronal function^4–7^. In the retinofugal system, studies in mice using optic nerve crush (ONC) have identified downregulation of mechanistic target of rapamycin (mTOR) signaling as a major driver of dendritic loss after axonal damage^8^. Consistent with a neuroprotective role for mTOR in this context, preventing its suppression after ONC preserves dendritic architecture and improves injury outcomes^9^. Moreover, stimulating mTOR signaling with insulin can even promote dendritic regrowth after degeneration took place^10,11^.

Contrary to mammals, zebrafish possess a remarkable regenerative capacity. Following ONC, adult zebrafish do not undergo retinal ganglion cells (RGCs) loss and recover most vision-guided behaviors within approximately 2 weeks. Despite this resilience, the inner plexiform layer (IPL) of injured zebrafish undergoes an early but reversible thinning, reflecting transient dendritic pruning. Notably, IPL thickness is restored only after successful reinnervation of brain targets^12^. Disrupting this pruning response delays regeneration, suggesting that dendritic plasticity plays a permissive and potentially integral role in successful axonal regrowth^12^. Additional evidence from *C. elegans*^13^ and rodent models of induced axonal regeneration^14,15^ supports a role for dendritic shrinkage in conditioning injury-induced axonal regeneration. The spontaneously regenerating adult zebrafish therefore represents a powerful model to dissect dendritic remodeling during axonal regeneration and to identify paradigms with translational potential for mammalian systems. Here, we use a transgenic zebrafish line, the *Tg(gfi1ab:gal4;UAS:lynGFP)*^*16*^, with sparsely labelled RGCs to define the morphological features of dendritic pruning and regrowth during axonal regeneration and to identify the molecular mechanisms that drive this process.

With quantitative analysis of RGC dendritic morphology in *Tg(gfi1ab:gal4;UAS:lynGFP)* fish^16^, we demonstrate that axonal injury and regrowth are associated with transient dendritic simplification. The pruning phenotype reached its maximum extent by 6 days post-injury (dpi) and was primarily characterized by a loss of branching and a concurrent decrease in total dendritic length. Dendritic architecture then recovered as axons re-established tectal innervation, approaching uninjured complexity by 21 dpi. This structural remodeling program was accompanied by a biphasic mTOR response: mTOR activity increased acutely after injury, but components of the PI3K–Akt– mTOR cascade were subsequently downregulated at both the transcript and post-translational level during the dendritic pruning and axonal elongation phase. Enhancing insulin signaling upstream of mTOR, or stimulating mTOR directly, partially preserved dendritic complexity but delayed optic tectum reinnervation. These findings delineate a mechanistic link between transient dendritic pruning and efficient axonal regrowth, and may inform therapeutic strategies to improve late stages of optic nerve and CNS repair in mammals at large.

## Results

### Characterization of gfi1ab-expressing RGCs in the adult zebrafish retina

Characterization of RGC dendritic dynamics at single cell resolution requires sparse cell labelling. In this study, we used transgenic *Tg(gfi1ab:gal4;UAS:lynGFP)* zebrafish^16^, in which Gal4 expressed in *gfi1ab*+ cells drives the UAS-mediated expression of a membrane-bound GFP reporter (Fig. 1A-B). Similarly to what has been shown in larvae^16^, this transgenic approach results in sparse labeling of RGCs predominantly in the central to mid-peripheral dorsal retina of adult zebrafish (Fig. 1B). The labelled RGCs innervate the stratum fibrosum griseum superficiale (SFGS), stratum griseum centrale (SGC) and stratum album centrale/stratum periventriculare (SAC/SPV) of the dorsomedial optic tectum of adult fish (Fig. S1A-E). Overall, this pattern of labelling allows for imaging of isolated RGCs in retinal wholemounts and the ability to perform single-cell morphometric analyses of dendritic complexity and stratification in the inner plexiform layer (IPL) (Fig. 1B-C).

**Figure 1.**
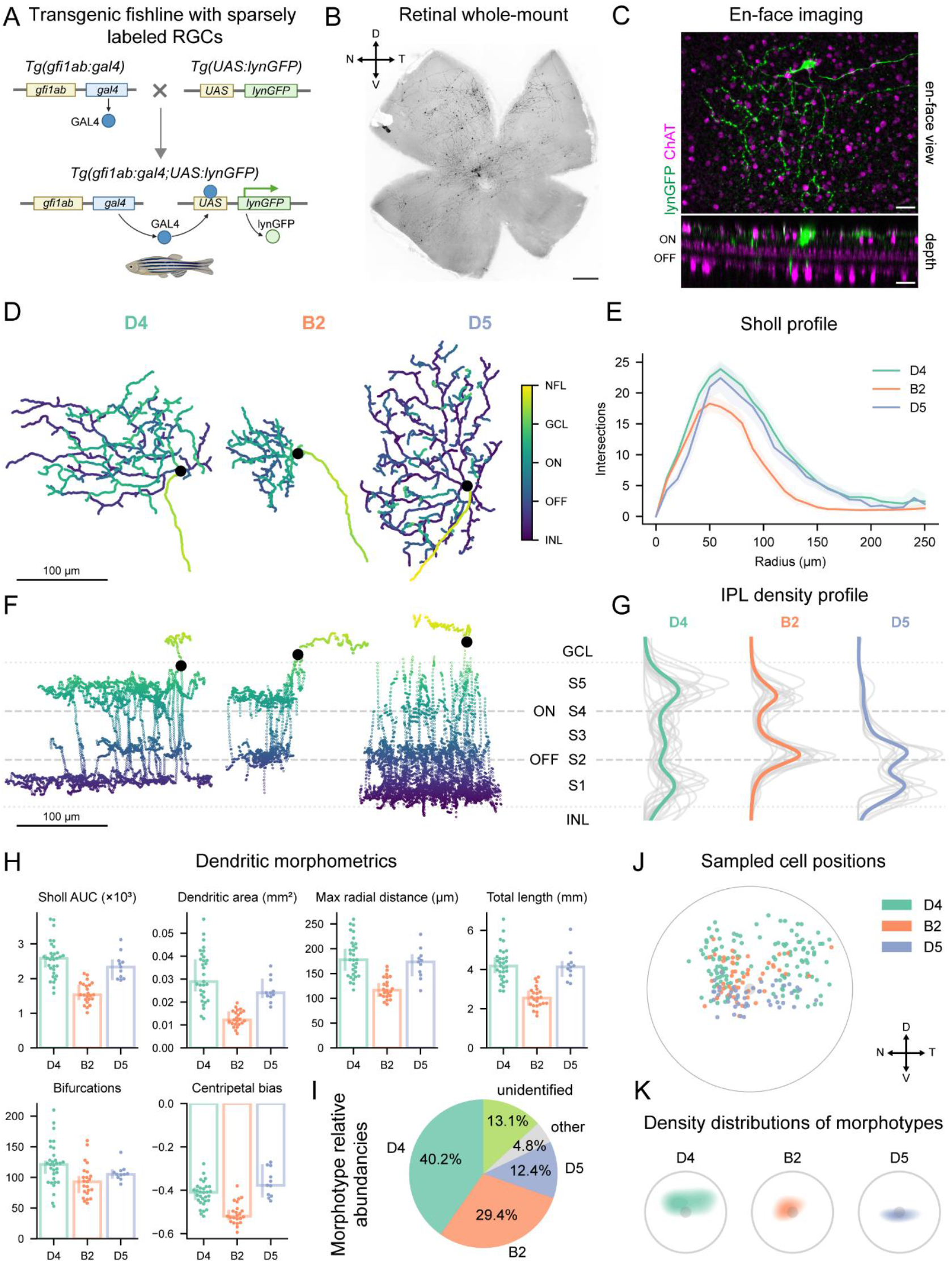
gfi1ab-driven sparse labelling reveals distinct adult zebrafish RGC morphotypes and IPL stratification patterns. (A) Schematic representation of the *Tg(gfi1ab:Gal4;UAS:lynGFP)* transgenic zebrafish line. Gal4 expression is regulated by the *gfi1ab* promoter and drives expression of a membrane-bound GFP (lynGFP) reporter via the *UAS* promoter. (B) Representative retinal whole-mount showing sparsely labeled RGCs, enriched in central and mid-peripheral regions of the dorsonasal retina. Scale bar = 250 µm. (C) En-face view (top) and optical section (bottom) of representative immunostaining for lynGFP (gfi1ab-driven) and choline acetyltransferase (ChAT), labeling starburst amacrines, identifies stereotyped IPL bands used to normalize dendritic depth measurements and minimize mounting-related distortion. Scale bar = 20 µm. (D) En-face (XY) dendritic reconstructions of representative D4, B2, and D5 morphotypes; D4 and D5 cells display larger, more elaborate arbors than B2 neurons. Color indicates relative position along the inner plexiform layer, with purple close to the inner nuclear layer (INL) and yellow close to the nerve fiber layer (NFL). Scale bar = 100 µm. (E) Sholl profiles reveal bigger and more complex dendritic arbors for D4 and D5 relative to B2 cells. (F) Orthogonal (XZ) reconstruction of dendritic stratification of D4, B2, and D5 morphotypes within the IPL. Sublaminae are defined relative to ON and OFF ChAT bands (see Fig. S2). Color is defined as in panel D. (G) Depth profiles across the IPL indicate a bistratified B2 pattern (S2 and S4), whereas D4 and D5 cells show broader, diffuse stratification, with D5 dendrites skewed toward OFF sublaminae. (H) Morphometric summary across arbor size and complexity metrics highlights similarity between D4 and D5 neurons and lower values for B2 cells. (I) Relative abundancy of the identified morphotypes across all sampled cells sampled. Rare morphotypes are grouped as “other”, while cells that could not be identified due to low-confidence ChAT band registration are labelled as “unidentified”. (J) Spatial coordinates of D4, B3 and D5 RGCs sampled across the retina. (K) Density distribution of sampling locations by morphotype, showing dorsal enrichment of D4, modest dorso-nasal enrichment of B2, and ventro-central enrichment of D5 neurons. Abbreviations: Area under the curve (AUC); choline acetyltransferase (ChAT); dorsal (D); inner plexiform layer (IPL); nasal (N); retinal ganglion cell (RGC); Sublamina (S); temporal (T); ventral (V).

To characterize the cells labeled by this line, we manually traced the dendrites of individual RGC and measured their stratification within the inner plexiform layer (IPL) using choline acetyltransferase (ChAT) staining as a reference (Fig. 1C, Fig. S2A-C, see Methods). Although the diversity of RGC morphological types has not been described in depth in the adult zebrafish retina^17^, larval RGCs have been more extensively classified^18^. We identified three main RGC morphological types: two similar to known larval types and one that appears to be novel (Fig. 1D,F). The two known types correspond to larval morphotypes D4 and B2: both exhibit directionally biased dendritic trees but differ in dendritic field size and lamination (Fig. 1D–H). D4 cells have larger trees and stratify mainly in ON band S5 and OFF band S1, with additional branching in S3 (Fig. 1D–H). B2 cells, by contrast, stratify only in ON band S5 and OFF band S2 (Fig. 1D–H). The novel type, which we named D5 expanding the establish nomenclature of larval RGCs with diffuse stratification^18^, has a large, complex dendritic tree that stratifies predominantly in OFF bands S1 and S2, with some extensions into the ON bands (Fig. 1D–H). D4 cells are the most abundant (∼40%), followed by B2 (∼30%) and D5 (∼15%) (Fig. 1I). Other morphotypes previously identified in the larval retina, such as B1 and B5 (Fig. S2D), were occasionally observed, but their low frequency (<2% for both) prevented a full quantitative analysis of their dendrites. Despite our biased sampling towards sufficiently isolated cells, we observed a robust and diverse spatial distribution for the three main morphotypes. D4 cells were mainly found in the dorsal retina, with a slight bias toward the dorsonasal quadrant (Fig. 1J,K). B2 cells also localized to the dorsal retina, with a stronger dorsonasal bias (Fig. 1J,K). In contrast, D5 cells were located on the ventral side of the central retina, proximal to the optic nerve head (Fig. 1J,K). Overall, the *Tg(gfi1ab:gal4;UAS:lynGFP)* line is well suited to analyze dendritic complexity at the single-RGC level, and labeling is largely restricted to three distinct morphological RGC types at the adult stage.

### A reduction of dendritic complexity after optic nerve crush precedes brain reinnervation

To determine how dendrites remodel during axon regeneration, we quantified the dendritic architecture of RGCs in uninjured retinas and at several timepoints post-ONC, chosen to capture the key stages of regrowth based on previous studies^12,19–21^. These stages included: 2 dpi, corresponding to the end of the initiation phase of axonal regrowth, 3 dpi, when most axons are extending within the optic nerve and tract; 6 dpi, when axons begin to reinnervate the tectum; 10 dpi, when tectal reinnervation is largely complete but vision remains impaired; 14 dpi, when basic visual behaviors are restored; and 21 dpi, when complex vision is nearly fully recovered (Fig. 2A). To confirm that this overall regeneration timeline (Fig. 2A) is representative for *gfi1ab*+ axons specifically, we traced regenerated axons in optic tectum whole-mounts at 3, 6, 10 and 14 dpi (Fig. S1F). Representative traces revealed only rare pioneering axons entering the tectum 3 dpi. By 6 dpi, axons had entered the tectum from the nasal and temporal sides, but most did not reach their target in the center of the tectum. From 10 dpi to 21 dpi the central tectum was repopulated by the terminals of the regenerated *gfi1ab*+ axons.

**Figure 2.**
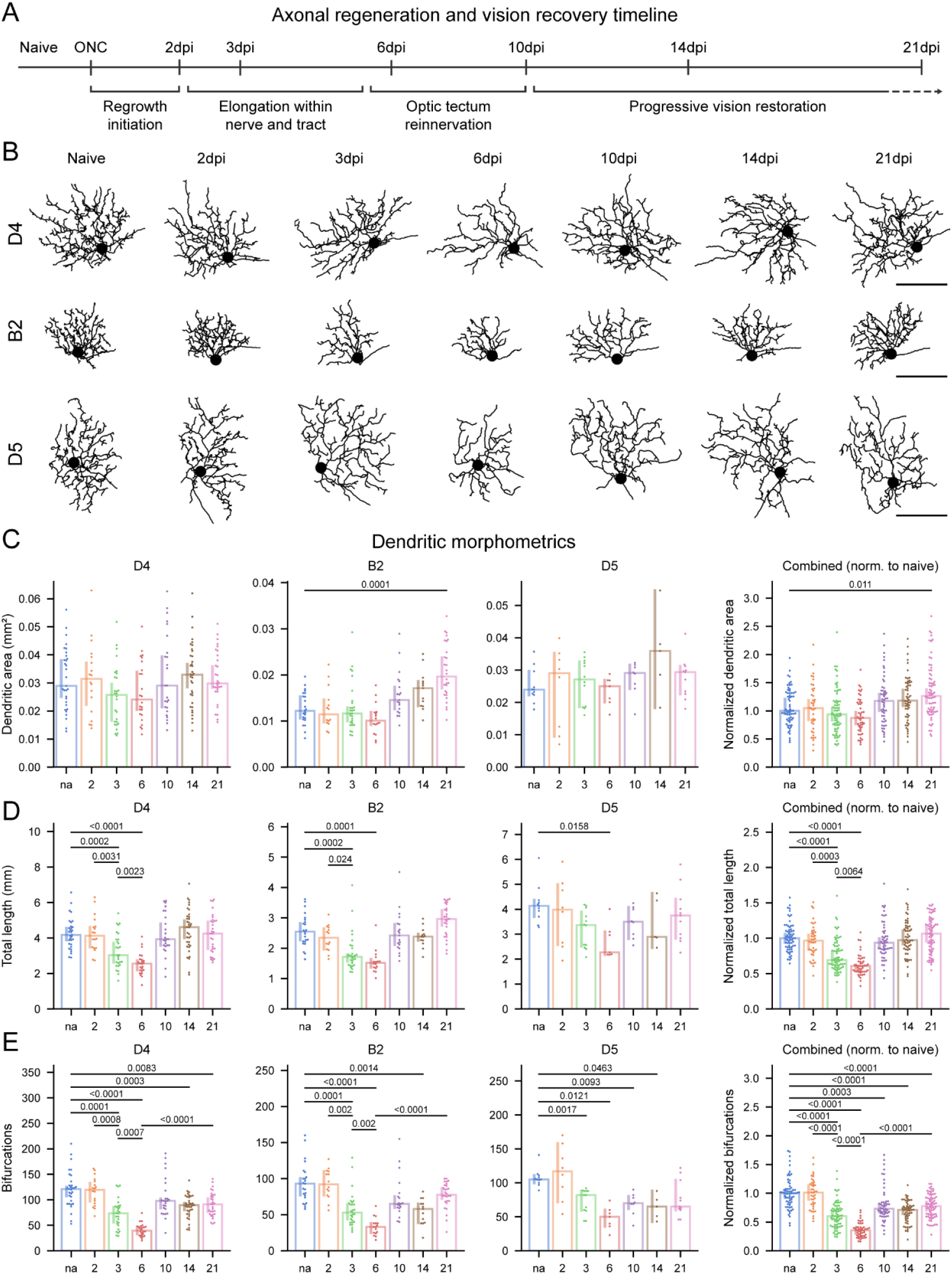
Optic nerve crush induces transient dendritic pruning that precedes tectal reinnervation. (A) Experimental timeline showing imaging/quantification time points relative to phases of axon regrowth, tectum reinnervation, and vision recovery. Representative traces of *gfi1ab*+ axons in the tectum from 3–14 days post-injury (dpi) are shown in Fig. S1. (B) Dendritic reconstructions of representative D4, B2, and D5 RGCs across time points after optic nerve crush, illustrating reduced arbor complexity at 3 and 6 dpi and restoration after 10dpi. Scale bar = 100 µm. (C–E) Quantification of dendritic area (C), total dendritic length (D), and number of bifurcations (E) for each morphotype across regeneration reveals significantly reduced dendritic complexity, but not decreased field area, at 3dpi and 6dpi relative to naive and 21dpi. Values are shown for all morphotypes individually as well as combined, after normalization to the mean of the corresponding uninjured morphotype. Data presented as median ±95% confidence interval. Each dot represents a cell (N=40-70 cells sampled across n=10 fish per timepoint). One-way Kruskal-Wallis ANOVA with multiple comparisons followed by Mann–Whitney post hoc tests; p values are shown in the figure. Abbreviations: days post-injury (dpi); optic nerve crush (ONC).

ONC did not affect dendritic complexity at 2 dpi but induced a marked reduction thereafter (Fig. 2B–E), with ∼30% decreases in total bifurcation number, total dendritic length (Fig. 2D–E), and the area under the curve (AUC) of the Sholl profile (Fig. S3A–B) across all three morphotypes at 3 dpi. Dendritic simplification progressed further between 3 and 6 dpi, particularly in the total number of bifurcations, which declined to ∼60% of uninjured levels (Fig. 2D–E). In contrast, the overall dendritic field area was not significantly altered (Fig. 2C), suggesting that remodeling primarily involves the elimination of terminal branches (Fig. 2A). By 10 dpi, total dendritic length and Sholl profiles were restored in all three morphotypes (Fig. 2D, Fig. S3A–B). In contrast, although the total number of bifurcations showed a significant recovery at 21 dpi compared with 6 dpi, it did not reach uninjured levels within the time window examined (Fig. 2E). This suggests that fine-tuning of distal dendritic segments and their associated synaptic contacts might continue beyond this timepoint. Across morphotypes, dendritic remodeling dynamics were largely similar over the regeneration timeline. An exception was morphotype B2, which exhibited a significant increase in dendritic arbor area at 21 dpi, potentially reflecting ongoing late-stage dendritic finetuning. Collectively, these findings indicate that a transient, highly regulated dendritic simplification phase accompanies axon regrowth and that dendritic morphology is largely re-established upon full target reinnervation in the tectum, before recovery of visual function.

### A transient downregulation in expression of the insulin cascade underlies dendritic remodeling and axonal regeneration

To gain insight into the molecular players underlying the dendritic remodeling, we made use of a recently published bulk RNA sequencing dataset of isolated adult zebrafish RGCs after ONC (Fig. 3A)^22^. Gene set enrichment analysis (GSEA) on the Gene Ontology (GO) cellular component dataset revealed a significant downregulation of terms associated with pre- and post-synapses, starting from 1 dpi and reaching the maximum extent between 3 dpi and 6 dpi (Fig. 3B). Terms associated with dendrites were also significantly downregulated, most prominently at 3 dpi and 6 dpi (Fig. 3B). GSEA on the GO Biological Process and Reactome datasets showed a similar downregulation of transcripts associated with dendrite development, synaptic assembly and postsynaptic transmission between 1 dpi and 6 dpi, and most prominently at 3dpi (Fig. 3C).

**Figure 3.**
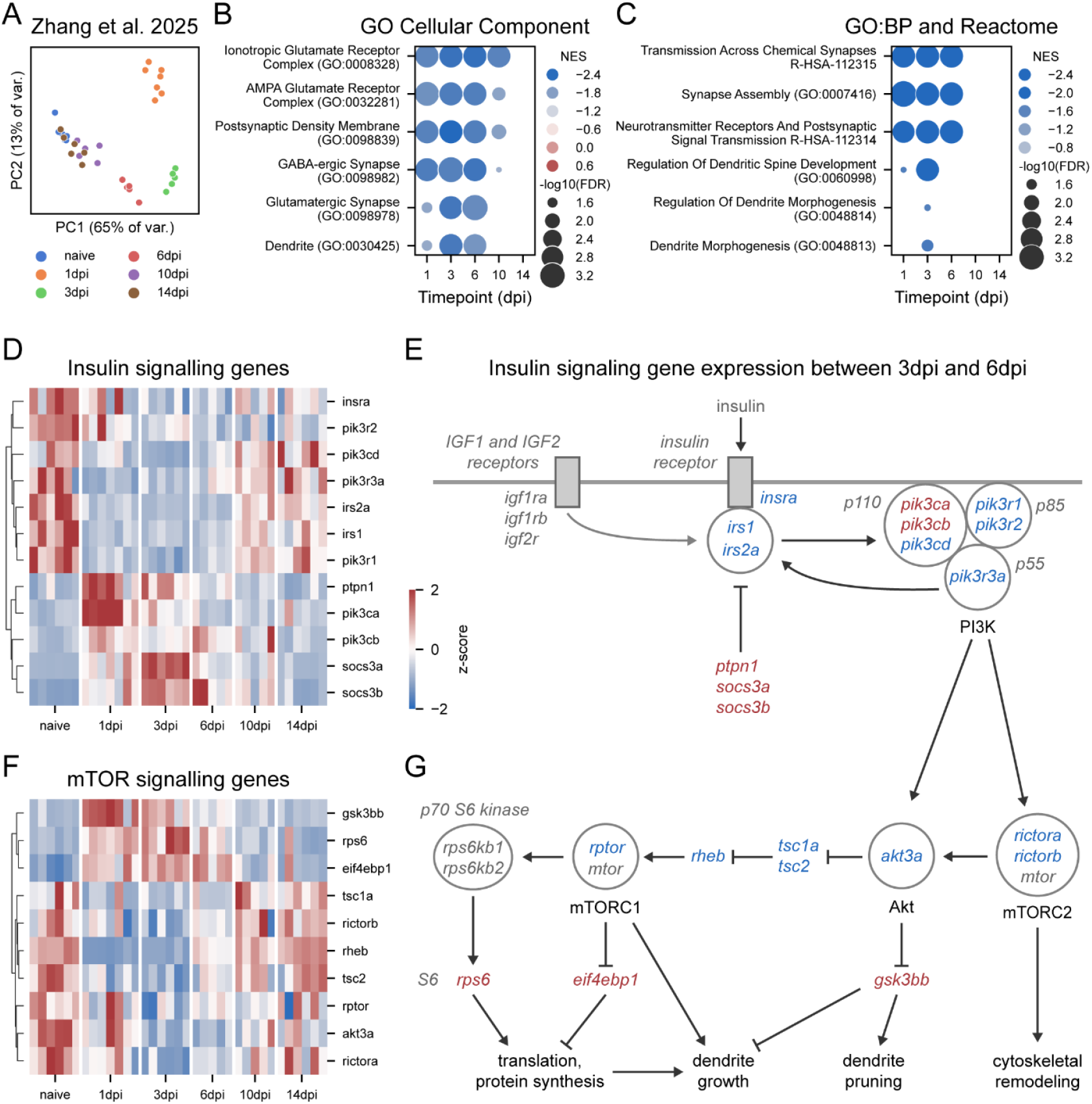
Transient transcriptional suppression of insulin signaling coincides with dendritic remodeling and axonal regeneration. (A) Principal component analysis (PCA) of the Zhang et al., 2025 dataset containing transcriptomes of retinal ganglion cells (RGCs) from uninjured control (naive) fish and from fish subjected to ONC at 1, 2, 3, 6, 10, and 14 days post-injury (dpi). (B-C) Dot plot of GSEA on the GO Cellular component, GO Biological Process and Reactome databases. Synaptic and dendritic-related terms are predominantly downregulated between 1 and 6 dpi. Terms that were upregulated are marked in red, and those downregulated in blue. (D-E) Heatmap and schematical representation of genes involved in insulin signaling, showing expression pattern consistent with downregulated signaling between 1 and 6 dpi. Genes that are upregulated are marked in red, and those downregulated in blue. (F-G) Heatmap and schematic representation of mTOR pathway genes downstream of insulin signaling, showing decreased expression of major pathway components between 1 and 6 dpi. Genes that are upregulated are marked in red, and those that are downregulated in blue. Heatmaps show per-sample z-scores across time points. Genes with similar expression dynamics are grouped hierarchically. Abbreviations: days post-injury (dpi); Gene Ontology (GO); False Discovery Rate (FDR); gene set enrichment analysis (GSEA); mechanistic target of rapamycin (mTOR); Normalized Enrichment Score (NES); principal component analysis (PCA); retinal ganglion cells (RGCs).

We next asked whether insulin–mTOR signaling participates in dendritic remodeling, given its established roles in supporting dendrite stability and growth in RGC stress paradigms (e.g., glaucoma) as well as across CNS development and degeneration^8,23^. Consistent with the synaptic/dendritic gene sets, the insulin–PI3K–Akt–mTOR cascade exhibited coordinated downregulation at the transcriptional level after injury, with suppression apparent by 1 dpi, maximal at 3 dpi, and sustained through 6 dpi (Fig. 3D–G). At the pathway entry point, the receptor *insra* was modestly downregulated at 3 and 6 dpi (log2FC −1.02 and −1.06), while *igf1ra*/*igf1rb*/*igf2r* were unchanged; notably, the initiating adaptor nodes *irs1* and *irs2a* were reduced from 1–6 dpi (Fig. 3D–E). Downstream, regulatory PI3K subunits (*pik3r1*/*2*/*3*) decreased over the same time window, accompanied by isoform-specific alterations in catalytic subunits (increased α/β, decreased δ), and a concomitant rise in negative regulators of IRS signaling (*ptpn1, socs3a/b*) (Fig. 3D–E). Further along the cascade, key components spanning mTOR complexes and their control nodes (*akt3a, rptor, rictora*/i, *rheb, tsc1a*/*tsc2*) were reduced at 3–6 dpi, whereas translation-related genes (*rps6, eif4ebp*) were elevated throughout 1–6 dpi (Fig. 3F–G), suggesting potential decoupling between the upstream pathway and its downstream effectors during the remodeling phase. In parallel, *gsk3bb*—an Akt-regulated kinase linked to dendrite pruning^24^—was strongly induced after injury (Fig. 3F–G).

Together, these changes point to a transient reduction in trophic signaling that likely weakens synaptic and dendritic maintenance programs during the period of maximal axonal elongation. Consistent with this model, expression patterns largely normalized by 10 dpi, when tectal reinnervation is complete and dendritic structure is mostly restored.

### Injury-induced dendritic pruning is associated with repressed mTOR signaling rather than Gsk3β activity

Given the extensive post-translational control of mTOR signaling, RNA expression alone provides an incomplete proxy for pathway activity. We therefore assessed mTOR pathway activation in situ via immunohistochemistry for pS6 (Ser235/236), an in-tissue proxy readout for mTORC1 activation, and for pGsk3β (Ser9), which reflects Akt-dependent inhibitory phosphorylation of Gsk3β (Fig. 4A). In uninjured retinas, pS6-positive cells were rare in the ganglion cell layer (GCL). Following ONC, the fraction of pS6-positive GCL cells increased to ∼50% at 1–2 dpi, but rapidly declined to near-undetectable levels between 3 and 10 dpi (∼5%) (Fig. 4B-C). This defined temporal pattern suggests that mTORC1 activation is transient and is sharply repressed by 3 dpi, coinciding with the onset of pronounced dendritic pruning.

**Figure 4.**
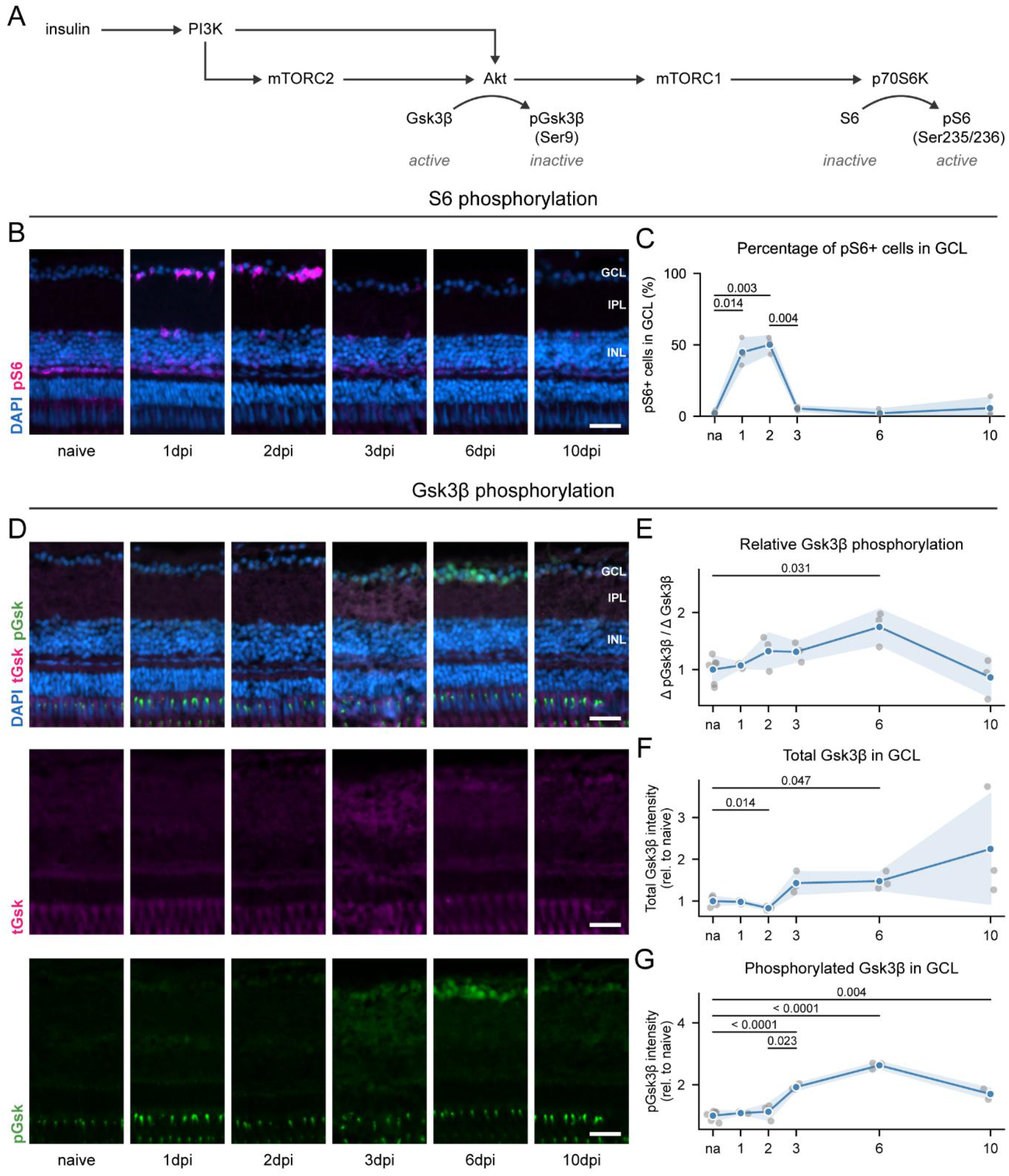
Dendritic pruning following ONC is associated with decreased mTOR signaling rather than increased Gsk3β activity. (A) Schematic representation of insulin signaling, showing AKT-dependent inhibitory phosphorylation of GSK3β (Ser9) and mTORC1/p70S6K-dependent phosphorylation of ribosomal protein S6 (Ser235/Ser236) (pS6). (B) Representative sagittal retinal cryosections stained for phospho-S6 (pS6, Ser235/Ser236) and DAPI show increased S6 phosphorylation at 1 and 2 dpi. Scale bar = 25 µm. (C) Quantification of pS6–positive cells in the ganglion cell layer (GCL) in naive retinas and at indicated days post-injury (dpi) reveals transient elevated mTORC1 activity at 1–2 dpi. (D) Representative sagittal retinal cryosections immunolabeled for total GSK3β, phospho-GSK3β (Ser9), and DAPI show an increase in total GSK3β and phospho-GSK3β (Ser9) from 3 to 10 dpi, peaking at 6 dpi. Scale bar = 25 µm. (E-G) Quantification of fluorescence intensity for total GSK3β and phospho-GSK3β (Ser9), and the phospho/total ratio reveal that total GSK3β and Ser9 phosphorylation increase at 3–6 dpi. Data presented as mean ± SD (C-G). Each dot represents one fish (n=3-5). One-way Welch ANOVA with Games–Howell multiple-comparisons test; exact p values are shown in the figure. Abbreviations: days post-injury (dpi); ganglion cell layer (GCL); inner plexiform layer (IPL); inner nuclear layer (INL), mechanistic target of rapamycin (mTOR); optic nerve crush (ONC); mechanistic target of rapamycin (mTOR); phospho-GSK3β (Ser9) (GSK3β); phospho-S6 (Ser235/Ser236) (pS6).

Considering the elevated expression of Gsk3β and its potential role in destabilizing microtubules and actin networks during dendritic pruning, we next examined its phosphorylation status (Fig. 4D-C). Total Gsk3β levels increased significantly at 6 dpi (Fig. 4D,F), whereas inhibitory Ser9 phosphorylation was elevated as early as 3 dpi and remained high through 10 dpi compared with uninjured retinas (Fig. 4D,G). Consequently, the Ser9 phosphorylation ratio was higher at 6 dpi (Fig. 4E), indicating a net shift toward inhibitory modification and reduced Gsk3β activity despite increased protein abundance. Overall, these data support a model in which dendritic pruning between 3 and 6 dpi is associated primarily with suppression of mTORC1 signaling rather than enhanced Gsk3β activity.

### Prolonged insulin signaling activation results in reduced dendritic remodeling and impaired brain reinnervation

To determine whether reduced insulin/mTOR pathway activity contributes to injury-induced dendritic pruning, we exogenously sustained signaling beyond 2 dpi by intravitreal administration of human recombinant insulin. This strategy has been shown to preserve and restore dendritic arbors in murine glaucoma models through mTOR-dependent mechanisms^10,11^. Insulin was injected after ONC, at 2, 3, and 4 dpi (Fig. 5A) and elicited an expected downstream signalling response, increasing the percentage of pS6+ cells in the GCL ∼3-fold at 3 dpi compared with vehicle (Fig. 5B), despite the identified moderate transcriptional downregulation of insulin receptors (Fig. 3D, E). Although dendritic complexity declined from 3 to 6 dpi in both conditions (Fig. 5C–D), insulin-treated retinae exhibited significant arbor preservation at 6 dpi, with increased Sholl AUC, dendritic area, total dendritic length, and branch points relative to vehicle (Fig. 5C–D). Similar trends were observed across all morphotypes individually (Fig. S4A). These data indicate that insulin treatment induces partial preservation of RGC dendritic architecture following ONC injury.

**Figure 5.**
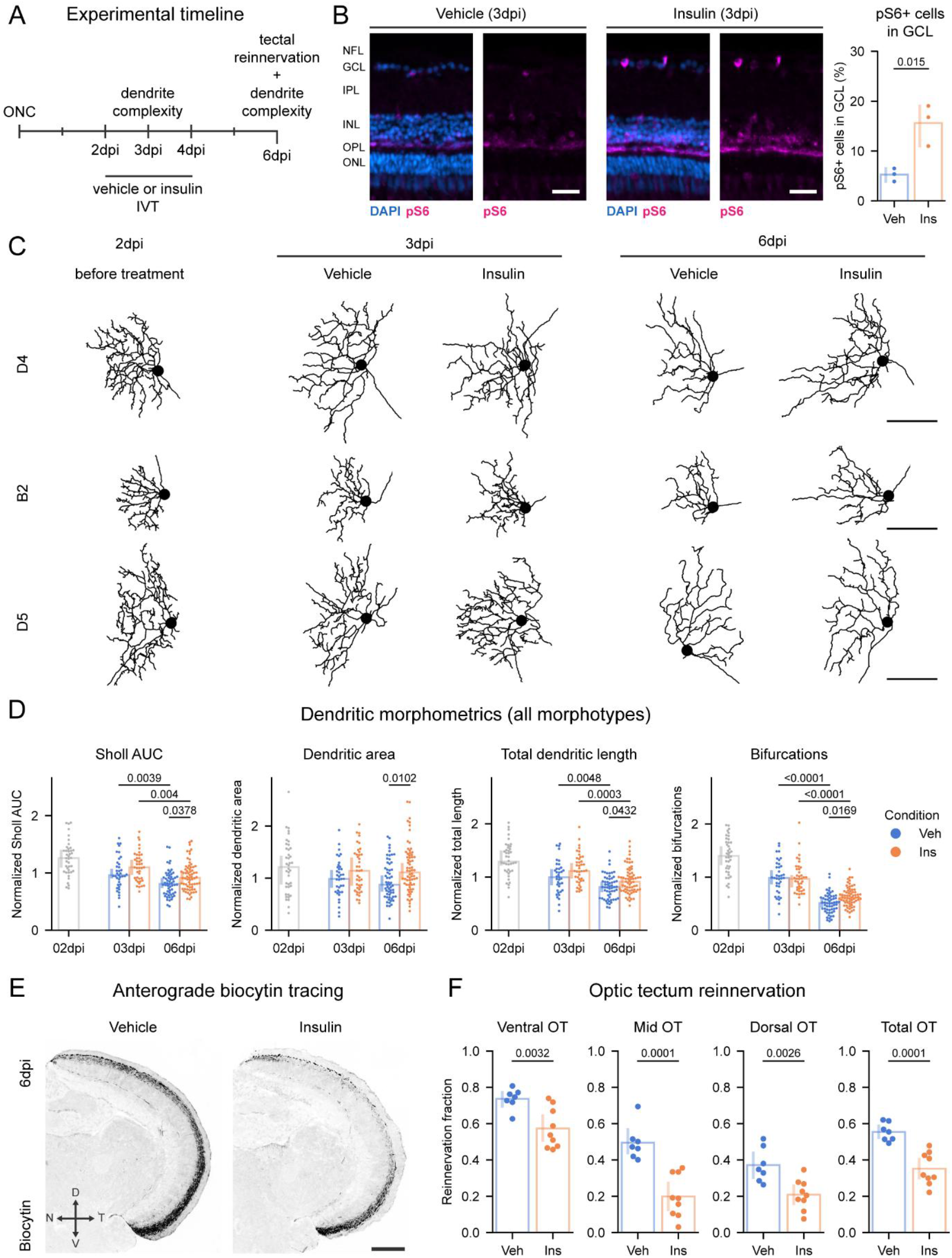
Sustained insulin stimulation following ONC results in reduced dendritic pruning and impaired brain reinnervation. (A) Experimental timeline showing vehicle (PBS) or recombinant human insulin (0.2 mg/mL) intravitreal injections at 2, 3, and 4 days post-injury (dpi). Dendritic morphology was quantified at 3 and 6 dpi, and optic tectum reinnervation was assessed at 6 dpi to evaluate delays in axonal regrowth. (B) Representative sagittal retinal cryosections stained for phospho-S6 (pS6, Ser235/Ser236) and DAPI; quantification of pS6–positive cells in the ganglion cell layer (GCL) reveals that insulin administration increased the fraction of pS6+ cells at 3dpi, consistent with enhanced mTORC1 activation. Scale bar = 25 µm. (C) Representative dendritic reconstructions following ONC at 2 dpi (pre-treatment) and at 3 and 6 dpi after vehicle or insulin treatment show increased dendritic complexity of RGCs in insulin-treated retinas compared with vehicle-treated controls. The representative reconstructions at 2 dpi are taken from Fig.2 as a reference for the baseline before treatment. Scale bar = 100 µm. (D) Quantification of dendritic morphometrics (Sholl AUC, dendritic area, total dendritic length, and number of bifurcations) shows that both experimental groups exhibit pruning over time. Insulin-treated RGCs retain higher dendritic complexity at 6 dpi relative to vehicle-treated. The 2 dpi data (grey) are taken as reference from Fig. 2 for comparison with the insulin- and vehicle-treated groups. E) Representative images of coronal optic tectum sections after anterograde biocytin tracing. Biocytin visualization shows reduced tectal reinnervation at 6 dpi upon insulin treatment. Scale bar = 200 µm. (F) Quantification of the biocytin-positive area in the entire optic tectum and in ventral, medial, and dorsal regions reveals that insulin treatment delays tectal reinnervation compared with vehicle. Data presented as median ± 95% confidence interval (D) or mean ± SD (B,F). Each dot represents one cell (D; N=40-47 cells from n=5-8 fish at 3 dpi, N=71-74 cells from n = 10 fish at 6dpi) or one fish (F; n = 7–8 per group). Student t-test (B), one-way Kruskal-Wallis ANOVA with multiple comparisons followed by Mann–Whitney post hoc tests (D) or Student t-test (F); exact p values are shown in the figure. Abbreviations: area under the curve (AUC); days post-injury (dpi); dorsal (D); insulin (Ins); intravitreal injection (IVT); phosphorylated ribosomal protein S6 (pS6); ganglion cell layer (GCL); nasal (N); optic nerve crush (ONC); optic tectum (OT); standard deviation (SD); temporal (T); vehicle (Veh); ventral (V).

We next assessed axonal regeneration by quantifying optic tectum reinnervation at 6 dpi by anterograde biocytin tracing. Daily insulin treatment from 2–4 dpi reduced tectal reinnervation by ∼50% relative to vehicle, indicative of a delay in axonal regrowth (Fig. 5E–F). Together, these results demonstrate that early insulin administration post-injury partially preserves RGC dendritic complexity but substantially delays brain reinnervation, supporting a model in which a precisely timed, transient suppression of insulin/mTOR signaling promotes dendritic pruning and enables timely axon extension in regenerating adult zebrafish RGCs.

### Downregulation of mTORC1 signaling is required for sustained axonal regrowth

To define the relative contribution of downstream effectors of insulin signaling to dendritic pruning and axonal regeneration, we tested whether rapamycin —a specific mTORC1 inhibitor—could reverse the effects of insulin and whether direct pharmacological targeting of mTOR with MHY1485 —a small molecule designed to activate mTOR by targeting its ATP domain— could mimic the effects of insulin (Fig. 6A-C). Compounds were delivered by intravitreal injection, daily between 2–4 dpi (Fig. 6A-C). Combined treatment with insulin and rapamycin significantly reduced pS6 phosphorylation in the GCL compared with vehicle, confirming effective mTORC1 inhibition. In contrast, MHY1485 administration did not produce a detectable increase in pS6 levels relative to vehicle under these experimental conditions (Fig. 6C-C).

**Figure 6.**
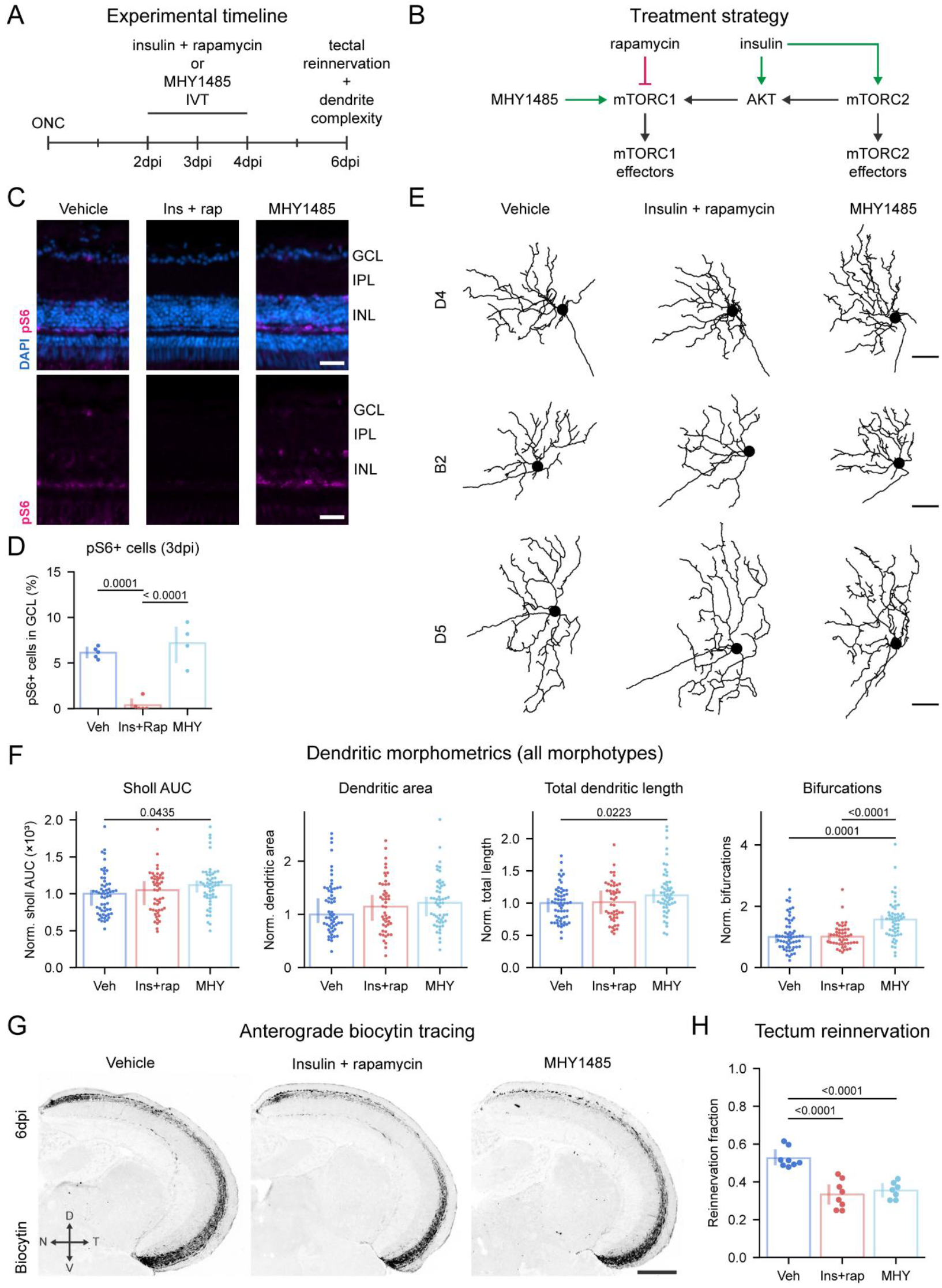
Downregulation of mTOR signaling is required for sustained axonal regrowth. (A) Experimental timeline showing intravitreal injections of vehicle (1% DMSO), insulin + rapamycin (0.2 mg/mL and 20 µM, respectively), or MHY1485 (100 µM) at 2, 3, and 4 days post-injury (dpi). Dendritic morphology and optic tectum reinnervation are quantified at 6 dpi. (B) Schematic of the insulin signaling pathway and pharmacological manipulations. Insulin activates mTORC2 and, via Akt signaling, mTORC1. Rapamycin selectively inhibits mTORC1, while MHY1485 selectively activates it. Green arrows indicate activation, and red blunt-ended lines indicate inhibition. (C) Representative sagittal retinal cryosections stained for phospho-S6 (pS6, Ser235/Ser236) and DAPI show comparable S6 phosphorylation in MHY1485-treated and vehicle-treated retinas at 3 dpi. Combined insulin and rapamycin treatment sharply reduces S6 phosphorylation in the retina. Scale bar = 25 µm. (D) Quantification of pS6–positive cells in the ganglion cell layer (GCL) shows a comparable percentage of pS6-positive cells in MHY1485-treated and vehicle-treated retinas, whereas combined insulin and rapamycin treatment results in almost no pS6-positive cells in the GCL. (E) Representative dendritic reconstructions at 6 dpi of cells from retinas treated with vehicle, combined insulin and rapamycin, or MHY1485. Combined insulin and rapamycin treatment results in dendritic complexity comparable to vehicle controls, whereas MHY1485-treated RGCs show increased dendritic complexity relative to vehicle-treated cells. Scale bar = 100 µm. (F) Quantification of dendritic morphometrics (Sholl AUC, dendritic area, total dendritic length, and number of bifurcations) reveals that MHY1485 increases dendritic complexity relative to vehicle, whereas insulin + rapamycin does not significantly alter dendritic complexity compared with vehicle. (G) Representative coronal optic tectum sections following anterograde biocytin tracing. Biocytin visualization shows decreased tectal reinnervation after both combined insulin and rapamycin treatment or MHY1485 treatment compared with vehicle controls at 6 dpi. Scale bar = 200 µm. (H) Quantification of biocytin-positive area in the optic tectum disclose that both insulin plus rapamycin and MHY1485 treatments delay tectal reinnervation relative to vehicle. Data presented as median ± 95% confidence interval (D) or mean ± SD (F). Each dot represents one cell (D; N=52-63 cells sampled from n = 7 fish per group) or one fish (D; n = 4-5) (F; n = 7). One-way Kruskal-Wallis ANOVA with Tukey HSD post hoc tests (D, H) or one-way Welch ANOVA with Games–Howell multiple-comparisons test (F); exact p values are shown in the figure. Abbreviations: area under the curve (AUC); days post-injury (dpi); dorsal (D); insulin (Ins); ganglion cell layer (GCL); nasal (N); rapamycin (Rap); phospho-S6 (pS6, Ser235/Ser236); temporal (T); vehicle (Veh); ventral (V).

At 6 dpi, rapamycin reversed insulin-dependent dendritic maintenance across morphotypes, restoring dendritic metrics to vehicle levels (Fig. 6E-C). Although pS6 levels were unchanged, MHY1485 induced a modest but significant increase in Sholl AUC and total dendritic length, and more than a 50% increase in bifurcations number compared with vehicle (Fig. 6E-C). Similar trends were observed when morphotypes were analyzed separately (Fig. S5A). Combined, these data suggest that the effect of insulin on dendritic preservation is mTORC1-dependent.

Assessment of axonal regeneration revealed that MHY1485 administration reduced tectal reinnervation at 6 dpi by ∼50% relative to vehicle, similar to what was measured following insulin treatment (Fig. 6 G-H). In contrast, although rapamycin fully reversed insulin-dependent dendritic maintenance, it did not restore tectal reinnervation, as animals treated with insulin plus rapamycin also exhibited ∼50% reduced reinnervation relative to vehicle-treated fish (Fig. 6 G-H). As mTOR signaling in RGCs is repressed only between 2 and 3 dpi in non-treated fish (Fig. 4 B-C), the observed effect may result from rapamycin prematurely inactivating mTORC1 at 2 dpi, thereby compromising the initiation of axonal regrowth in a dendrite-independent manner. Additionally, we cannot rule out the possibility that intravitreal rapamycin may affect the proximal optic nerve and acts directly on axons, contributing to reduced reinnervation. Overall, the data indicate that insulin-dependent dendritic maintenance is mTORC1-dependent, and that direct pharmacological inhibition of pruning through mTOR targeting is detrimental to axonal regeneration. These findings position transient dendritic pruning coupled to mTORC1 repression as a key driver of efficient axonal elongation and timely reinnervation of brain targets.

## Discussion

In this study, using quantitative measurements of single-cell morphology, we show that RGC dendrites undergo pruning during injury-induced axonal-regrowth in the spontaneously regenerating adult zebrafish, and that their architecture is restored upon brain reinnervation. This transient simplification phenotype coincides with the time window of axonal extension in the optic nerve, and is characterized by the loss of dendritic branching, while largely preserving the overall dendritic arbor area. Molecularly, we found that dendritic pruning is mediated by a temporary reduction in signaling of the insulin-mTOR axis. Interventions that sustain mTOR activation during axonal extension, either through direct pharmacological activation or by enhancing the upstream insulin signaling, attenuate dendritic pruning and severely impair optic tectum reinnervation. Therefore, mTOR-dependent maintenance of dendritic branching emerges as a limiting factor for sustained axonal regrowth and function recovery in the adult zebrafish.

### A defined set of gfi1ab+ RGC morphotypes is labelled in the adult zebrafish retina

Although the morphological heterogeneity of RGCs in the larval zebrafish is well established^18^, how this diversity is represented in the adult retina remains relatively unexplored. An earlier study described eleven morphological RGC subtypes in adulthood^17^, but a comprehensive classification is still lacking. In our study, we used the recently generated *Tg(gfi1ab:gal4;UAS:lynGFP)* zebrafish line, which labels a sparse set of various RGC morphotypes in the larval retina^16^. Based on dendritic morphology, we identified 3 major RGC morphotypes labelled by *gfi1ab* expression in the adult retina. Two of these appear to correspond to the larval D4 and B2 RGC subtypes described by Robles et al.^18^, and resemble subtypes IX and X reported by Mangrum et al.^17^. The third one, D5, appears to possess a novel stratification pattern that is diffuse across the IPL, but heavily skewed toward the OFF laminae. Whether this represents a truly novel morphotype or results from refinement of larval stratification during maturation into adulthood remains unclear. Longitudinal studies following larval RGCs as they develop into adult stages could help clarify this question. Regarding projections in the brain, in the adult optic tecum, *Tg(gfi1ab:gal4;UAS:lynGFP)* fish display 3 main clusters of layering: in the SFGS, SGC and SAC/SPV, respectively.

As different tectal layers process largely distinct stimuli, this hints at a potential difference in the functional properties between the three types. However, the precise projection patterns and functional characteristics of these adult RGC morphotypes remain largely unknown, and their detailed characterization lies beyond the scope of this study, which focuses on dendritic remodeling upon ONC injury.

### Dendritic pruning is an early phenotype of axonal damage

Dendritic pruning has received substantial attention in vision research, as it constitutes an early degenerative response to axonal damage in both acute trauma/axotomy models^6,10,15^, and glaucomatous conditions^11,25–28^. In these contexts, dendritic shrinkage contributes to the functional decline of neurons and precedes cell loss. Consequently, extensive research efforts have been devoted to preserving dendritic morphology as a strategy to prevent RGC degeneration. However, dendritic pruning after ONC is not exclusively a degenerative hallmark but is also observed in regeneration-competent systems. Robust dendritic pruning has been observed during the early phases of axonal regeneration in adult zebrafish and in murine RGCs with *Pten* and *Socs3* co-deletion^15^. Similarly, evidence from *C. elegans*^13^ and other regeneration-competent rodent models^14^ suggest that maintaining dendritic arbors can constrain axonal regrowth. Supporting this view, during RGC development in both fish and mice, axons extend first, and dendritic arborization occurs only after the axons have innervated their target areas in the brain^29–31^. This sequential order may have evolved to avoid competition between axonal and dendritic processes and could also be important in an adult setting for successful functional recovery. From this perspective, while uncontrolled dendritic degeneration is likely detrimental from a neuroprotective standpoint, the occurrence of finely tuned dendritic pruning as part of pro-regenerative programs that recapitulate development sequences may create a permissive environment that facilitates axon extension.

### A multifaceted role for mTOR in injury-induced axonal regeneration

In this work, we show that transient dendritic pruning in adult zebrafish after ONC is associated with marked downregulation of mTOR signaling, and particularly of mTORC1 activity. Mechanistically, this mirrors observations in murine RGCs, where dendritic shrinkage after ONC or in the microbead model of glaucoma is likewise accompanied by reduced mTOR activity. In these models, preventing mTOR downregulation via the knockdown of the negative regulator REDD2 preserves dendritic architecture^9^, and enhancing insulin signaling after the onset of dendritic shrinkage is sufficient to promote dendritic regrowth^10,11^. More broadly, mTOR functions as a master regulator of cellular growth that integrates extracellular growth factor signaling with intracellular nutrient and energy status to coordinate cap-dependent translation, ribosome biogenesis, metabolism, lipid synthesis and autophagy^23^. Activation of the PI3K–Akt–mTOR axis is a key determinant of neuronal survival and regenerative competence in the CNS^32,33^.

Accordingly, enhancing its activity through PTEN inhibition or related strategies is both necessary and sufficient to elicit robust axonal regrowth in injured mammalian CNS neurons^34–36^. Within this framework, the downregulation of mTOR observed in fully regenerating zebrafish RGCs might at first seem counterproductive. However, it suggests instead that precise temporal modulation of mTOR signaling is a prerequisite for successful regeneration. Supporting this view, we and others^12,21^ have measured a rapid and pronounced increase in mTOR activity in zebrafish RGCs during the first 2 days after injury, which likely serves to initiate the regenerative program. Blocking this early activation with the selective mTORC1 inhibitor rapamycin severely impairs axonal regrowth, confirming the importance of mTOR signaling at this stage^12,21^. Subsequently, from 3 days post-injury onward, mTOR signaling is strongly and rapidly suppressed in zebrafish, both transcriptionally and post-translationally, coinciding with the onset of dendritic pruning. This later downregulation appears to be equally critical, as we found that experimentally prolonging the window of mTOR activation, either by boosting insulin signaling upstream or directly targeting of mTOR, diminishes dendritic pruning and severely delays optic tectum reinnervation. In terms of direct mTORC1 targeting, MHY1485 at the concentrations used here has been reported to activate mTORC1 via increased phosphorylation of the mTOR subunit^37,38^ and, in some cases, elevating S6 phosphorylation in retina of mice and of zebrafish larvae^39,40^. However, other studies have not observed any detectable increase in pS6, suggesting that MHY1485-induced signaling downstream of mTORC1 could be context- and model-dependent^38,41^. Consistent with this, in our system MHY1485 did not appreciably increase pS6 at 3 dpi, yet it phenocopied the rapamycin-sensitive effect of insulin in preserving dendritic complexity. These findings provide compelling evidence that the protective effect of insulin is mTORC1-dependent and raise the possibility that S6 phosphorylation is not required for this specific effect. Additional mechanistic studies will be necessary to directly test this and to pinpoint which mTOR outputs, such as autophagy regulation, mediate the effects described here. Collectively, our results support a biphasic model in which mTOR activation is essential for the initial injury response and the initiation of axonal regeneration, whereas its subsequent rapid downregulation drives dendritic pruning and is required for timely axonal elongation and target reinnervation.

### Intraneuronal resource allocation and compartmentalized signaling might underlie successful axonal regeneration

A growing body of evidence suggests that the limited regenerative capacity of adult mammalian CNS neurons may reflect an inability to redistribute growth-supporting resources to injured axons. In mature CNS neurons, mitochondria become stably positioned near synapses and do not remobilize after injury^42^, whereas regenerating zebrafish RGCs transiently relocate their mitochondria from dendrites into elongating axons^12^. Rab11-positive recycling endosomes show a similar maturation-dependent compartmentalization: they support growth in developing axons and growth cones but are largely confined to the somatodendritic domain in mature neurons, restricting regrowth unless axonal transport is restored^43,44^. Together, these findings support a competition model in which the maintenance of postsynaptic structures and dendritic arbors limits the pool of resources available for axon regeneration. Insulin–mTOR signaling could contribute to this framework, as insulin receptors are normally localized to the postsynaptic density^45–47^ and mTOR is enriched at postsynaptic sites within dendrites^48^, where it regulates local protein synthesis.

We found that intravitreal insulin delivery starting at 2 dpi activated the pathway but did not fully sustain mTOR signaling or completely prevent dendritic pruning. These findings align with the observed transcriptional downregulation of insulin receptors and several downstream components of the cascade, suggesting a reduced sensitivity of RGCs to insulin during axonal growth. On the other hand, signaling through the related IGF1 receptor at the growth cone is essential for axonal regeneration in hippocampal neurons^49^, and viral-mediated IGF1 overexpression promotes robust corticospinal tract regeneration^50^. Downstream of insulin, local PI3K signaling supports axonal regeneration of DRG neurons^35^ and mTOR has been shown to localize to growth cones^51–53^. Taken together, this evidence raises the compelling hypothesis that insulin signaling, and its downstream mTOR activity, become compartmentalized after injury in regenerating neurons. We propose that transient dendritic pruning represents an adaptive response, reallocating resources, such as protein synthesis capacity, membrane biogenesis, and energy metabolism away from dendrites to prioritize axonal regeneration.

### Limitations of the study and future perspectives

Our manuscript provides compelling evidence that temporally controlled dendritic remodeling is an integral component of successful injury-induced axonal regeneration in adult zebrafish. Although dendritic morphology was assessed in a limited set of RGC morphotypes, immunostainings for mTOR activity (or its absence) revealed consistent patterns across the entire retina, and pharmacological manipulations affected tectal reinnervation globally. Together these observations suggest that our findings are likely representative of the broader RGC population. Pharmacological manipulation was chosen because of the limited availability of reliable delivery tools, such as viral vectors, which render time- and spatially controlled interventions at the genetic level extremely challenging in adult zebrafish. Consequently, our study does not dissect the precise molecular mechanisms through targeted genetic perturbations.

Although we provide insights into how dendritic remodeling contributes to successful axonal regeneration in adult zebrafish, it remains unclear how this remodeling during axonal regrowth affects RGC electrical activity. Pruning of post-synaptic sites and dendritic branches is expected to alter the synaptic input that RGCs receive from bipolar and amacrine cells. At the same time, synapse elimination may be selective, enabling a transient rewiring of retinal circuitry that preserves neuronal activity during axonal regrowth. Supporting this idea, visual stimulation has been shown to promote axonal regrowth in murine RGCs, suggesting that activity-dependent mechanisms remain operative^54^. Accordingly, a detailed electrophysiological characterization of RGC function during this remodeling phase will be essential to define its functional consequences and to inform future therapeutic strategies.

Finally, mechanistic insight into potential compartmentalized molecular mechanisms in axons versus dendrites could be obtained using *in vitro* systems, such as microfluidic devices, which enable selective manipulation of individual neuronal compartments. While initial axonal regrowth can be induced in mammals, robust and extensive reinnervation of central brain targets remains largely unachieved, with only modest reinnervation observed in the most successful studies. Hyperactivation of mTOR signaling is a common feature of most of these interventions, yet our findings suggest that sustained mTOR activity might hinder the later phases of axonal regeneration. Exploring strategies that modulate dendritic remodeling together with precise spatiotemporal control of mTOR activation and inhibition could provide valuable avenues for promoting late-stage axonal regeneration in the mammalian CNS.

## Acknowledgments

We thank Evelien Herinckx and Arnold Van Den Eynde for the animal caretaking, Edzhem Akpanar and Astrid De Vriese for technical support. We thank Julie De Schutter, Pieter-Jan Serneels, Bram Nuttin, Dr. Joana Santos and Prof. Karl Farrow for the critical discussions. This research was funded by the FWO (project G082221N) and the KU Leuven Research Council (C14/22/074), by the IHU FOReSIGHT (ANR-18-IAHU-0001) (S.A. and F.D.B.)

European Research Council (ERC-2022-SYG grant 101071583 “TUBULINCODE”) (F.D.B.).

## Author contributions

Conceptualization: AZ, LuM, SB, LM; Data curation: AZ, LuM; Formal analysis: AZ, LuM; Funding acquisition: AZ, LuM, SB, LM; Investigation: AZ; Methodology: AZ, LuM, SB, EP, SA, LM; Project administration: AZ, LuM, LM; Resources: AZ, LM, FEP, SA, FDB, LM; Software: AZ, LuM; Validation: AZ, LuM, SB; Visualization: AZ, LuM, LM; Writing—original draft: AZ, LuM; Writing—review and editing: SB, EP, SA, FEP, FDB, LM.

## Declaration of interests

The authors have no actual or potential conflicts of interest.

## Resource availability

### Lead contact

Further information and requests for resources and reagents should be directed to and will be fulfilled by the lead contact, Lieve Moons

### Materials availability

All non-commercial reagents or fish lines used in this paper are available from the lead contact upon request. The *Tg(gfi1ab:gal4; UAS:lynGFP; cmlc2:eGFP)* fish line used in this study was obtained from the Del Bene lab (Institut de la Vision, Paris) and is available upon request.

### Data and code availability

- All original data reported in this paper will be shared by the lead contact upon request.
- The code used to train the ChAT detection deep-learning model is available at https://gitlab.com/NCDRlab/chat-train while the code used to normalize the dendritic coordinates in Z-stacks is available at https://gitlab.com/NCDRlab/stack_corr.
- Any additional information required to reanalyse the data reported in this paper is available from the lead contact upon request.

## Experimental model and study participant details

### Zebrafish maintenance

Zebrafish (*Danio rerio*) were maintained in a standardized flow-through system at 28 °C, with 15 fish per 3.5 L tank (ZebTec), under a 14 h light/10 h dark cycle. Fish were fed twice daily, with dry food in the morning and brine shrimp in the afternoon. All experiments were conducted on adult fish aged 5–9 months of both sexes and of similar size, as previous research has shown that the regenerative capacity of retinal RGCs is not affected by sex^55,56^. All procedures were approved by the KU Leuven Animal Ethics Committee and carried out in strict accordance with the European Communities Council Directive of 20 October 2010 (2010/63/EU).

We crossed *Tg(gfi1ab:gal4)* fish with the *Tg(UAS:lynGFP;cmlc2:eGFP)* line^16^. Offspring positive for a green fluorescent heart (driven by *cmlc2:eGFP*) and sparse GFP labeling on the yolk sac at 1 day post-fertilization were selected and raised to adulthood for experiments. In these *Tg(gfi1ab:gal4;UAS:lynGFP;cmlc2:eGFP)* fish, Gal4 is driven by the *gfi1ab* (Growth Factor Independent 1ab) promoter, which is active in a subset of RGCs^16,57^. Binding of Gal4 to the upstream activating sequence (UAS) induces expression of a membrane-localized GFP (lynGFP), enabling labeling of RGC neurites in a small subset of the RGC population. For experiments evaluating mTOR and Gsk3β activity on retinal cryosections, AB wild-type fish were used. The number of fish used per experiment is indicated by individual data points in the figure graphs and specified in the corresponding figure legends.

## Method details

### In vivo procedures

#### Optic nerve crush injury and tissue harvest

The unilateral optic nerve crush (ONC) injury was performed as previously described^55^. Briefly, fish were anesthetized with tris-buffered 0.03% tricaine (MS-222, MilliporeSigma, Burlington, MA, USA) solution in system water. The cornea surrounding the left eye was carefully removed using forceps. The eye was then gently lifted from the socket to expose the optic nerve. The optic nerve, approximately 0.5 mm from the optic nerve head, was crushed with Dumont #5 tweezers (Fine Science Tools, FST, Heidelberg, Germany) for 10 seconds. Afterwards, the eye was repositioned, and the fish was returned to fresh system water for recovery.

Under non-injured conditions or at 2, 3, 6, 10, 14, and 21 days post injury (dpi), fish were cardially perfused with PBS followed by 4% PFA. Eyes and brains were dissected and further fixed in 4% PFA (1 hour for eyes, overnight for brains). The eyes and optic tectum were subsequently wholemounted (Fig. S1C) and stained.

### Intravitreal injection and anterograde tracing of axonal regrowth

To augment insulin signaling during the axonal elongation phase, recombinant human insulin was injected intravitreally at 2, 3, and 4 days post ONC. Briefly, fish were anesthetized with buffered 0.03% tricaine in system water. The left eye was injected with 300 nL of 0.2 mg/mL recombinant human insulin (Sigma-Aldrich, Cat# I2643) using a 2.5 μL Hamilton syringe fitted with a 34G needle. Vehicle control fish underwent ONC and intravitreal injection at the same timepoints but received sterile PBS as a vehicle.

To enhance mTOR signaling activation, the mTOR activator MHY1485 (MCE, Cat# HY-B0795)^37^, was administered by intravitreal injection at 2, 3, and 4 dpi using the same procedure described above, at a final injection concentration of 100 µM. Vehicle control fish underwent ONC followed by intravitreal injection of 1% DMSO in sterile PBS in at the same timepoints. To verify that insulin signaling acts through mTORC1, insulin (0.2 mg/mL) was co-injected intravitreally with rapamycin (20 µM, MCE, Cat# HY-10219)^58^ at 2, 3, and 4 dpi using the same procedure.

To evaluate tectal reinnervation at 6 dpi, anterograde biocytin tracing was performed as reported previously^55^. Briefly, fish were anesthetized and prepared for surgery as describe above. The optic nerve was then transected between the nerve head and the crush site, and a piece of gelfoam soaked in biocytin was placed on the nerve stump to allow anterograde transport of biocytin into regenerating axons. After three hours, fish were cardially perfused, and brains were dissected and sectioned.

### Sample preparation for visualizing RGC dendrites and axonal reinnervation

Retinal wholemounts were stained for lynGFP to visualize *gfi1ab*-positive RGC dendrites and for ChAT (choline acetyltransferase, a marker of cholinergic starburst amacrine cells) to visualize the ChAT bands in the inner plexiform layer for stratification. Briefly, retinal wholemounts underwent a freeze–thaw cycle (15 minutes at –80 °C followed by thawing at room temperature) in 0.5% PBST (0.5% Triton X-100 in PBS). The samples were then incubated in blocking solution (2% PBST with 20% pre-immune donkey serum [PID, Sigma-Aldrich, Cat# S30-M]) overnight at room temperature. Subsequently, retinal wholemounts were incubated with primary antibody solution (2% PBST with 10% PID) containing chicken anti-GFP (1:500, Abcam, Cat# ab13970) and goat anti-ChAT (1:100, Sigma-Aldrich, Cat# AB144P) overnight at room temperature. After three washes, the wholemounts were incubated with secondary antibody solution (2% PBST with 10% PID) containing donkey anti-chicken Alexa Fluor 488 (1:200, Jackson, Cat# 703-545-155) and donkey anti-goat Alexa Fluor 647 (1:200, Invitrogen, Cat# A-21447) for 2 hours at room temperature. Following three additional washes, nuclei were stained with DAPI (1:1000 in PBS) for 2 hours. After a final PBS wash, tissues were mounted with Mowiol® using spacers between the carrier slide and coverslip avoid mechanical distortion of the dendrites.

Optic tectum wholemounts were stained for lynGFP to visualize gfi1ab-positive axons. As for retinal wholemounts staining, tectal wholemounts underwent a freeze–thaw cycle and blocking procedures as described above. Next, they were incubated with primary antibody solution (2% PBST with 10% PID) containing chicken anti-GFP (1:500, Abcam, Cat# ab13970) overnight at room temperature. After three washes, the samples were incubated with secondary antibody solution (2% PBST with 10% PID) containing donkey anti-chicken Alexa Fluor 647 (1:200, Jackson, Cat# 703-605-155) for 2 hours at room temperature. Nuclei were subsequently stained with DAPI (1:1000 in PBS) for 2 hours, and the tissues were mounted with Mowiol®.

Following perfusion and tissue dissection as detailed above, the eyes and brains were also prepared for cryosectioning. Samples were cryoprotected in a graded sucrose series (10%, 20%, and 30% sucrose in PBS), and samples were subsequently embedded in 1.25% agarose prepared with 30% sucrose in PBS. Sagittal cryosections of the eye (10 µm) and coronal cryosections of the brain (10 µm) were collected on SuperFrost® Plus Slides (Epredia, Cat# J1800AMNZ) using a CryoStar NX70 (Epredia).

Retinal sections were stained for pS6 (the phosphorylated form of S6) as an in-tissue proxy of mTOR (mechanistic Target of Rapamycin) activity and for Gsk3β and phospho-Gsk3β (Ser9). Briefly, sections were dried at 37 °C and rehydrated with deionized water. After washing with TBST (Tris-buffered saline containing 0.1% Tween-20), sections were incubated in blocking solution (20% PID in TBST) for 2 h at room temperature. Retinal sections were then incubated with primary antibodies diluted in antibody solution (10% PID in TBST) containing anti-pS6 (1:200; Cell Signaling Technology, Cat# 4857, RRID:AB_2181035) or containing anti-GSK3β (1:100; BioLegend, Cat# 604652) and phospho-GSK-3β (Ser9) (clone 5B3; 1:100; Cell Signaling Technology, Cat# 9323S; RRID:AB_2115201). Following three washes, sections were incubated with secondary antibody solution (10% PID in TBST) containing donkey anti-rabbit Alexa Fluor 647 (1:200, Invitrogen, Cat# A-31573) for pS6 staining or containing donkey anti-rabbit Alexa Fluor 647 (1:200, Invitrogen, Cat# A-31573) and donkey anti-mouse Alexa Fluor 555 (1:200, Invitrogen, Cat# A-31570) for GSK staining for 2 hours at room temperature. Following three additional washes, nuclei were counterstained with DAPI (1:1000 in PBS), and sections were mounted with Mowiol®. Brain sections were immunostained for lynGFP to visualize gfi1ab-positive axons and for biocytin to label RGC axons innervating the optic tectum. Sections were pretreated and blocked as described above. Next, they were incubated with primary antibody solution (10% PID in TBST) containing chicken anti-GFP (1:500, Abcam, Cat# ab13970) overnight. Following three washes, sections were incubated with secondary antibody solution (10% PID in TBST) containing donkey anti-chicken Alexa Fluor 647 (1:200, Jackson, Cat# 703-605-155) and streptavidin–Alexa 555 (1:300, Invitrogen, Cat# S32355) to visualize biocytin for 2 hours at room temperature. Following three additional washes, nuclei were counterstained with DAPI (1:1000 in PBS), and sections were mounted with Mowiol®.

### Bioinformatic analysis

#### Gene set enrichment analysis and heatmap generation

We reanalyzed a previously published bulk RNA sequencing dataset of FACS-sorted adult zebrafish RGCs under non-injured and injured conditions (1, 3, 6, 10, and 14 days post ONC injury)^22^. To identify pathways enriched within gene modules, we performed gene set enrichment analysis using GSEApy^59^ via the Enrichr API against the GO Cellular Component (GO:CC), GO Biological Process (GO:BP) and Reactome databases, using the DESeq2 Wald statistic (“stat”) as the ranking metric. Pathways with FDR *q* < 0.05 were considered differentially regulated. To visualize gene expression over regeneration time, differentially expressed genes were plotted with hierarchical clustering on z-scored heatmaps.

### Microscopic imaging

#### Imaging of single RGCs, optic tectum wholemounts, and brain cryosections

Retinal wholemounts were imaged using a Zeiss LSM900 confocal microscope equipped with a Plan-Apochromat 20×/0.8 NA objective. For imaging single RGCs, to capture the entire dendritic field, Z-stack images were acquired from the ganglion cell layer to the inner nuclear layer. For optic tectum wholemounts, Z-stacks covering the full tissue thickness were acquired and visualized as standard deviation projections. For brain cryosections, visualizing *gfi1ab*+ axons, Z-stacks encompassing the entire section thickness were acquired and visualized as maximum-intensity projections.

#### Imaging of retinal cryosections and optic tectum reinnervation

Images of retinal sections immunostained for pS6 and Gsk, as well as brain sections stained for biocytin following anterograde tracing, were acquired using a Leica DM6 microscope equipped with an HC PL FLUOTAR L 20×/0.40 CORR objective. Imaging settings were kept constant within each experimental batch.

### Image analysis and quantification

#### Tracing and quantification of RGC dendrites and axons

RGC dendrites and axons in confocal stacks were traced in Fiji/ImageJ using the SNT framework^60^. The traces generated for dendrites were then analyzed in Python with the NeuroM library^61^ to extract morphometric features of the dendritic arbors. The dendritic area was calculated by fitting a convex hull to the traced dendrites. Sholl area under the curve (AUC), dendritic area, total dendritic length, number bifurcations centripetal bias and maximum radial distance were chosen to define dendritic complexity. To analyze the complete RGC population regardless of morphotype, each cell was normalized to the mean of its morphotype control and normalized cells were then pooled.

#### Automated extraction of ChAT band positions and dendritic depth normalization for RGC morphotypes classification

To extract ON and OFF ChAT band positions, confocal 3D stacks were resampled into 5-µm XZ optical sections and sum-projected to generate 2D inputs. Manually annotated ON/OFF strata were used to train an attention U-Net for semantic segmentation of ChAT-positive bands. The model was optimized with a combined Tversky–Dice loss, adaptive learning-rate scheduling, and early stopping. Geometric deformation and intensity shift augmentations were applied to improve generalization. Segmentation accuracy was assessed using Dice coefficient and precision– recall metrics.

For inference, each section was evaluated with a sliding-window approach using overlapping tiles. Tile-level probability predictions were merged by averaging overlapping pixels to minimize edge effects, yielding dense probability maps for each lamina. Probability maps were thresholded to binary masks, spurious detections were removed by size-based filtering, and gaps in the resulting band traces were imputed by local linear interpolation. Traces were then smoothed with a Savitzky–Golay filter and mapped back to the corresponding Y coordinates in the original volume, providing ON and OFF band annotations throughout the stack.

Using the extracted ChAT bands as reference landmarks, dendritic reconstructions were normalized to a standardized IPL depth coordinate system following Sümbül et al.^62^. Low-confidence band annotations were removed and imputed by interpolation from neighboring valid sections. For each stack, smooth surfaces were fitted to the OFF and ON ChAT bands, and each dendritic point was assigned a normalized IPL depth defined as its fractional position between the two surfaces (OFF = 0, ON = 1). This transformation yields depth-normalized dendritic morphology, enabling quantitative comparisons across samples. After, normalized dendrite reconstruction was plotted as XY and XZ projection for manual morphotype classification. Morphotype identity was determined based on Robles et al.^18^. The D5 morphotype emerged as novel and was named as such to extend the classification of diffuse RGC morphology first presented by Robles et al.

#### Generation of density maps per morphotype

The sampling position of each cell was normalized to the position of the optic nerve head of the retina from which it was sampled, yielding radial coordinates. A 2D kernel density estimation was fit for each morphotype to plot its density relative to the optic nerve head.

#### Quantification of pS6 and Gsk3β

Immunostaining for pS6 and Gsk3β on retinal cryosections was quantified in QuPath. Nine equally spaced positions along the dorsal to ventral axis were imaged on a retinal cryosection per fish. On each image, regions of interest (ROIs) were manually drawn to include only the ganglion cell layer (primarily comprised of RGCs, but including also displaced amacrine cells). Nuclei were detected in QuPath using the DAPI channel, and a perinuclear cytoplasmic compartment was defined by expanding each nuclear object by 2 µm. For both pS6 and Gsk3β, mean fluorescence intensity was measured within the cytoplasmic compartment for each cell. For pS6, semi-automated intensity thresholding was used to classify pS6-positive cells on each image and to calculate their relative percentages. The percentages obtained from same retina were averaged to obtain a percentage of pS6-positive cells per fish. For Gsk3β, cytoplasmic intensities for phosphorylated Gsk3β (pGsk3β) and total Gsk3β were quantified per cell. For each marker, sample means were normalized to the mean of the naive condition. Relative Gsk3β phosphorylation was calculated as the pGsk3β/total Gsk3β ratio using the normalized intensities.

#### Quantification of optic tectum reinnervation

A custom ImageJ script was used to quantify tectal reinnervation, as described by Beckers et al.^55^. The area between the stratum opticum (SO) and the stratum fibrosum et griseum superficiale (SFGS) was manually delineated and measured as the total area of the SO and SFGS. The biocytin-positive area was then manually thresholded and measured. The percentage of the biocytin-positive area relative to the total SO/SFGS area was defined as the reinnervation percentage. Additionally, to measure the percentage of reinnervation in the dorsal, medial and ventral tectum, the total area was divided into three equally spaced region across the dorsal to ventral axis. For each fish, at least three sections of the central optic tectum were quantified, and the per-fish average reinnervation percentage of the analyzed sections was presented as one data point.

## Statistical analysis

Detailed information on the number of samples and biological replicates used are reported in the legends of the figures. For all experiments, the number of individual cells (N) and the number of fish (n) per condition or timepoint is indicated in the figure legends.

Raw, unsaturated micrographs were used for all quantitative analyses. For figure presentation only, images were occasionally inverted and/or contrast-adjusted by linear modification of the white point (or, for inverted images, the black point). Identical display adjustments were applied across images intended for direct comparison. For visualization of deep confocal z-stacks, rolling-ball background subtraction (ImageJ) was applied to facilitate display; background-subtracted images were not used for quantification. Details of statistical tests (e.g., t-tests and ANOVA) are provided in the figure legends. If normality was rejected in any group by the Shapiro–Wilk test, data are summarized as medians and analyzed using the Kruskal–Wallis test with pairwise Mann–Whitney post hoc comparisons. Otherwise, data are summarized as means and analyzed using Welch’s ANOVA with Games–Howell post hoc tests. All data processing, visualization, and statistical analyses were performed in Python using pandas, seaborn, matplotlib, statsmodels and SciPy. Statistical significance was defined as P < 0.05.

## Supplementary Material

**Figure S1.**
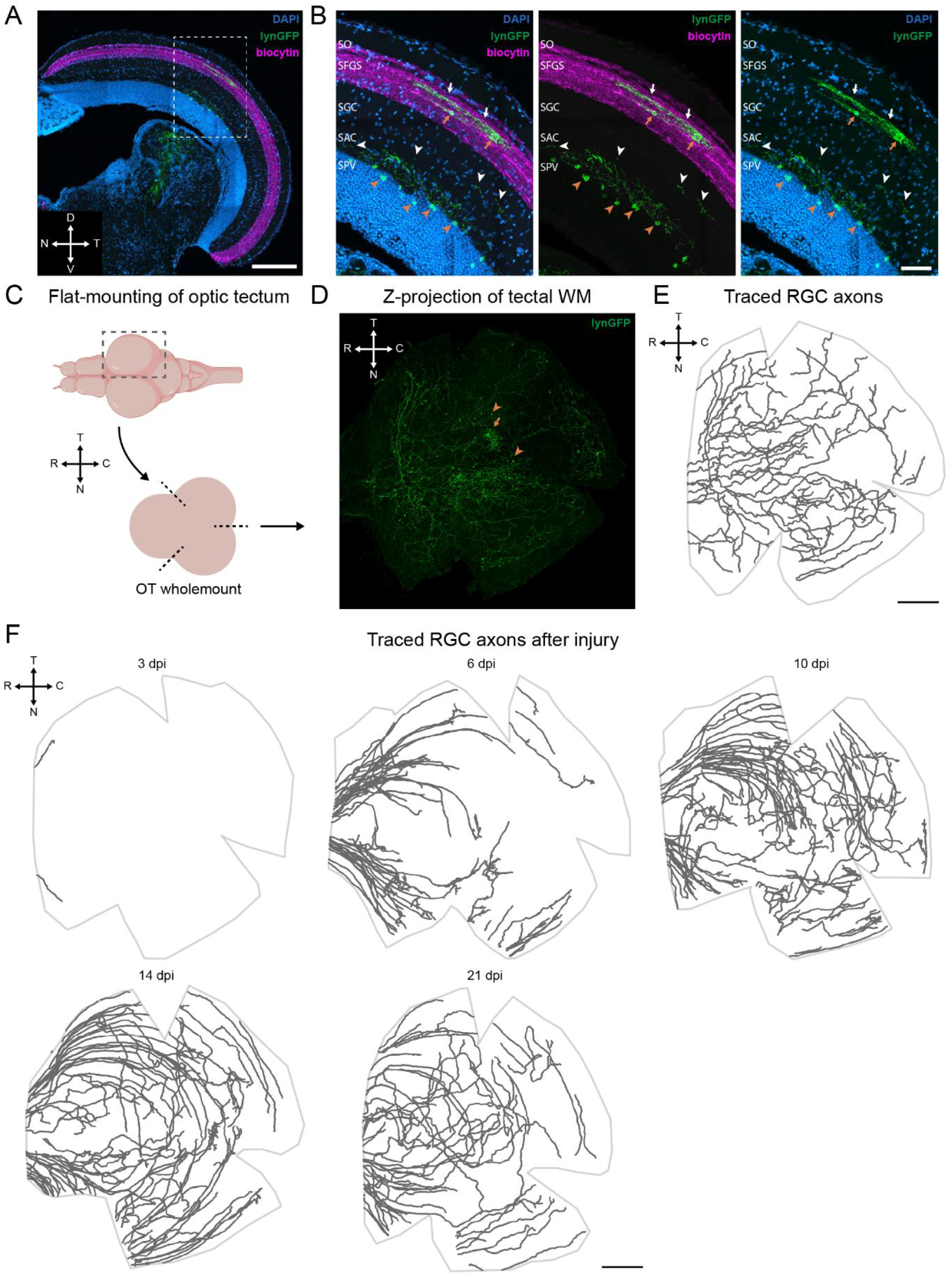
Tectal innervation pattern of gfi1ab+ RGCs and reinnervation timeline after injury. (A) Representative coronal brain cryosections from anterogradely traced fish labelled for DAPI, lynGFP and biocytin showing innervation of all RGCs (biocytin) and *gfi1ab*+ RGC axons in the dorsomedial optic tectum (white box) under non-injured conditions. Scale bar = 200 µm. (B) Magnified views of the coronal brain section reveal localized innervation by *gfi1ab*+ axons primarily in the SFGS (white arrows). Tectal neurons in the SFGS near the axon terminals are also sparsely labelled (orange arrows), possibly indicating synapses with the incoming RGC axons. Sparser innervation, mostly running perpendicular to the sectioning plane, is detectable also in the SGC and in the SAC (white arrowheads). Here, RGC axons appear in proximity with dendrites of labelled tectal neurons located in the SPV (orange arrowheads). See also panel D. Scale bar = 50 µm. (C) Schematic representation of optic tectum whole-mount preparation. The right optic tectum was peeled from the brain, followed by three cuts at the nasal–ventral, nasal–dorsal, and temporal positions. (D) Z-projection of a representative optic tectum wholemount showing innervation of *gfi1ab+* RGC axons to the medial optic tectum at non injured condition. Here, a small number of tectal neuron residing in the SGS are also labelled (orange arrow, see panel B). More abundant are neurons residing deeper in the SPV (orange arrowheads, see panel B). Scale bar = 200 µm. (E) Manually traced *gfi1ab+* RGC axons showing innervation in the central tectum at non injured condition. Scale bar = 200 µm. F) Representative reconstructions of manually traced *gfi1ab+* RGC axons showing the progressive reinnervation of the tectum after optic nerve injury. At 3 dpi, only rare pioneering *gfi1ab+* RGC axons are visible the optic tectum. By 6 dpi, a subset of labeled axons enters the optic tectum from the nasal and temporal sides and navigates toward targets in the central tectum. At 10 dpi, the majority of labeled axons navigate toward their target region. By 14 dpi, axons have reached the target in the central tecum; however, further refinement of innervation could take place after this timepoint. Scale bar = 200 µm. Abbreviations: caudal (C); choline acetyltransferase (ChAT); dorsal (D); days post-injury (dpi); inner plexiform layer (IPL); nasal (N); OT (optic tectum); retinal ganglion cell (RGC); rostral (R); temporal (T); ventral (V); whole-mount (WM).

**Figure S2.**
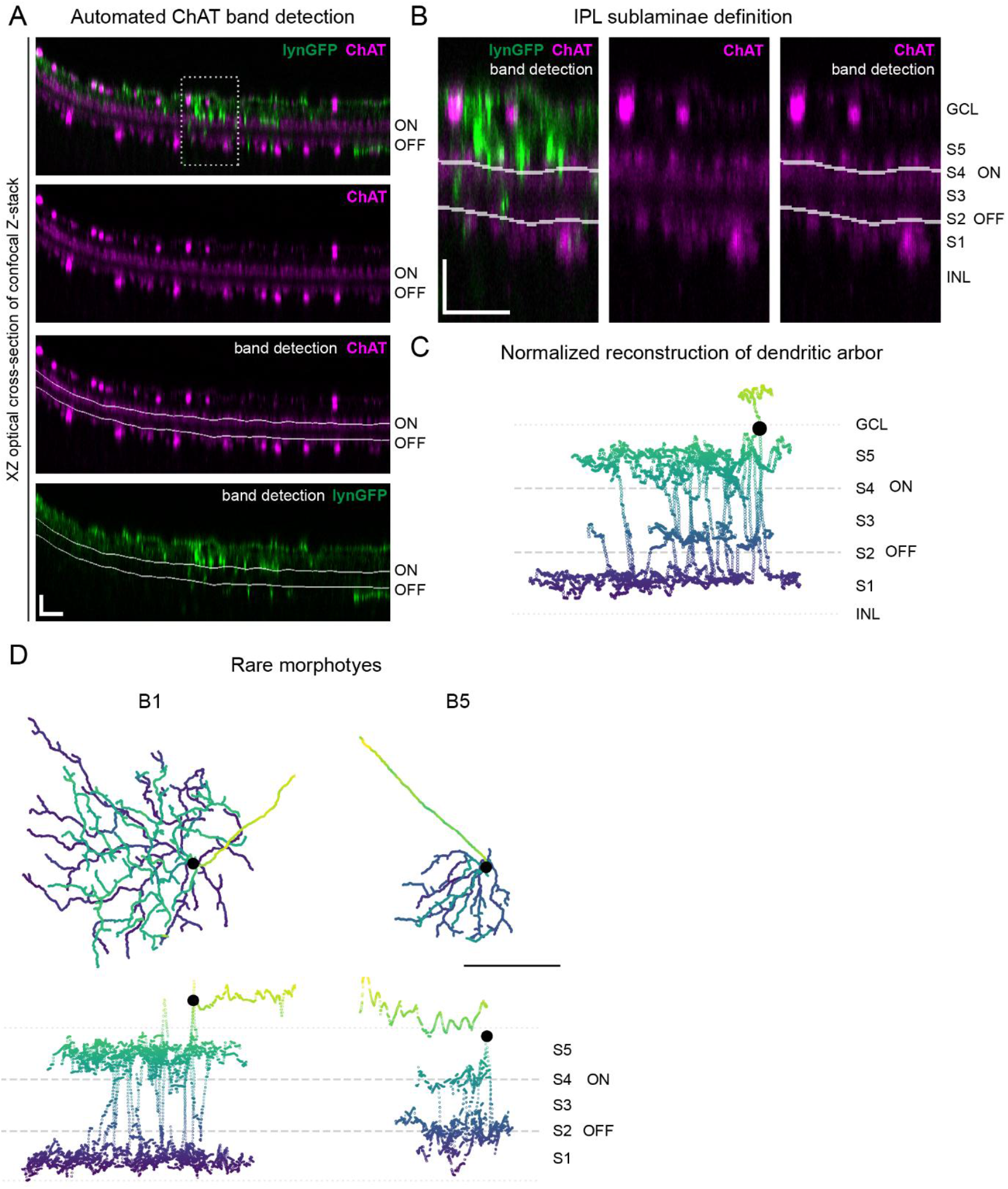
Normalization of RGC dendrite depth and representation of rare morphotypes. A) Optical section of a representative confocal stack used to image gfi1ab+ cells after immunostaining for lynGFP (green) and choline acetyltransferase (ChAT, magenta). An automated deep-learning-based model is run to detect the position of ON and OFF ChAT bands (white annotations) throughout the stack to be used as reference to normalize the coordinates of dendritic RGCs. This step is critical to cancel-out distortion artefacts due to irregular retinal whole-mounting. See Methods for a deeper methodological detail. Scale bars (vertical and horizontal) = 20 µm. B) Zoomed view of the optical section depicted in A. Chat band annotation are used to define IPL sublaminae, named S5 to S1 spanning from the GCL to the INL. S2 and S4 are defined in correspondence of the OFF and ON ChAT bands, respectively. Scale bars (vertical and horizontal) = 20 µm. C) After registration of coordinates relative to the reference ChAT band, normalized dendrite reconstructions are plotted. These faithfully represent the lamination of RGC dendrites within the IPL and are used to classify the 3 main morphotypes, namely D4, B2 and D5. D) En-face (XY, top) and orthogonal (XZ, bottom) dendritic reconstructions of representative rare B1 and B5 morphotypes. Although rarity prevents a quantitative characterization of their dendritic properties, B1 cells present large dendritic arbors stratified on S1 and S5, while B5 neurons present smaller arbors stratifying on S2 and S4. Scale bar = 100 µm. Abbreviations: choline acetyltransferase (ChAT); ganglion cell layer (GCL); inner plexiform layer (IPL); inner nuclear layer (INL).

**Figure S3.**
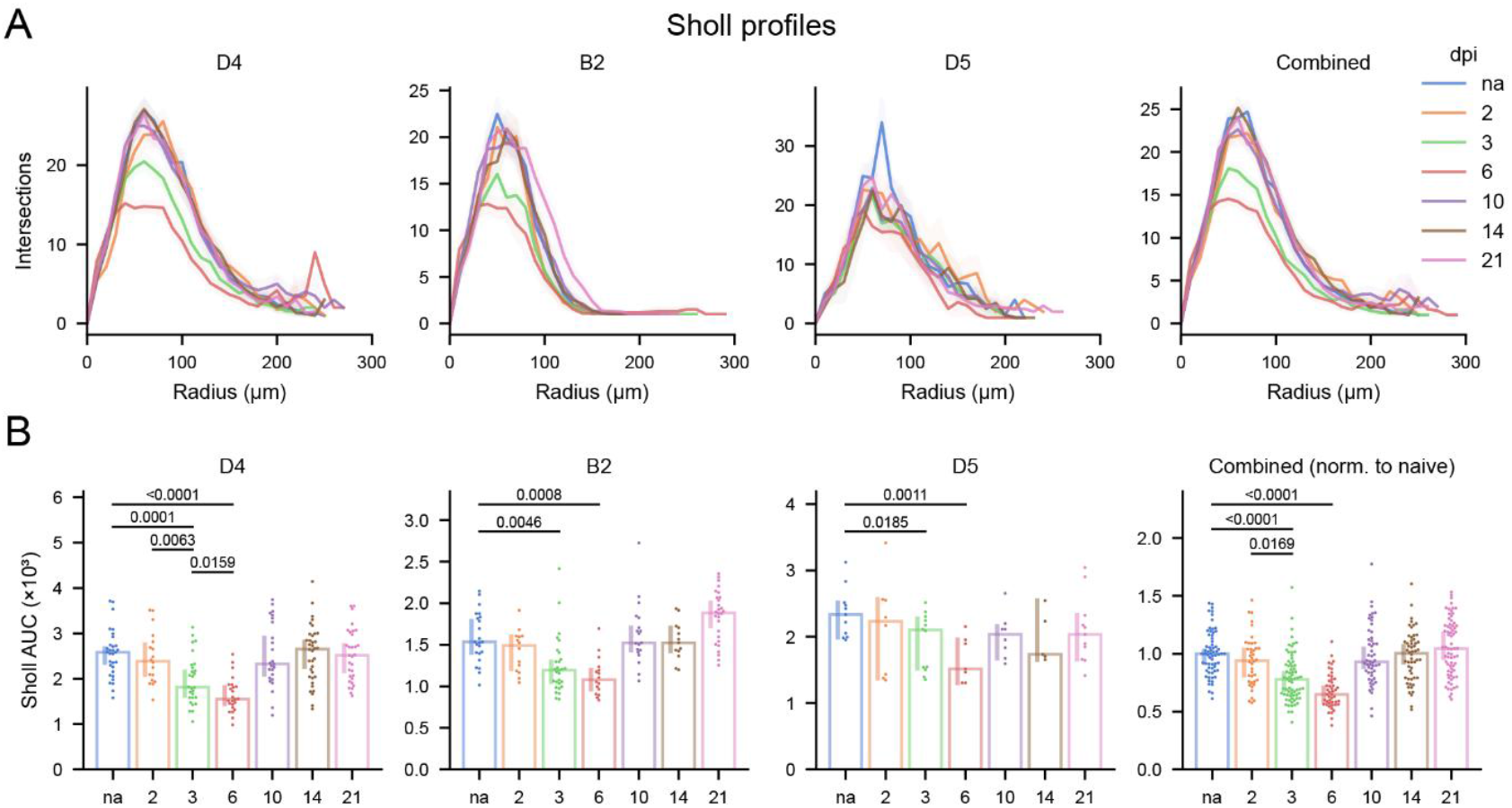
Sholl profiles of gfi1ab+ RGC morphotypes across the regeneration timeline. (A) Representative Sholl profiles of D4, B2, and D5 morphotypes, as well as the combined cell population, across the regeneration timeline. Following injury, all morphotypes exhibit a transient reduction in dendritic complexity, evident from day 3 (green line) and most pronounced at day 6 (red line). (B) Quantification of the area under the curve (AUC) of the Sholl profiles reveals significant dendritic pruning of RGCs at 3 and 6 dpi. Dendritic complexity is gradually restored by 10 dpi and further recovered through 21 dpi. Values are shown for all morphotypes individually as well as combined, after normalization to the mean of the corresponding uninjured morphotype Data presented as median ±95% confidence interval. Each dot represents a cell (N=40-70 cells sampled across n=10 fish per timepoint). One-way Kruskal-Wallis ANOVA with multiple comparisons followed by Mann–Whitney post hoc tests; p values are shown in the figure. Abbreviations: Area under the curve (AUC); days post-injury (dpi).

**Figure S4.**
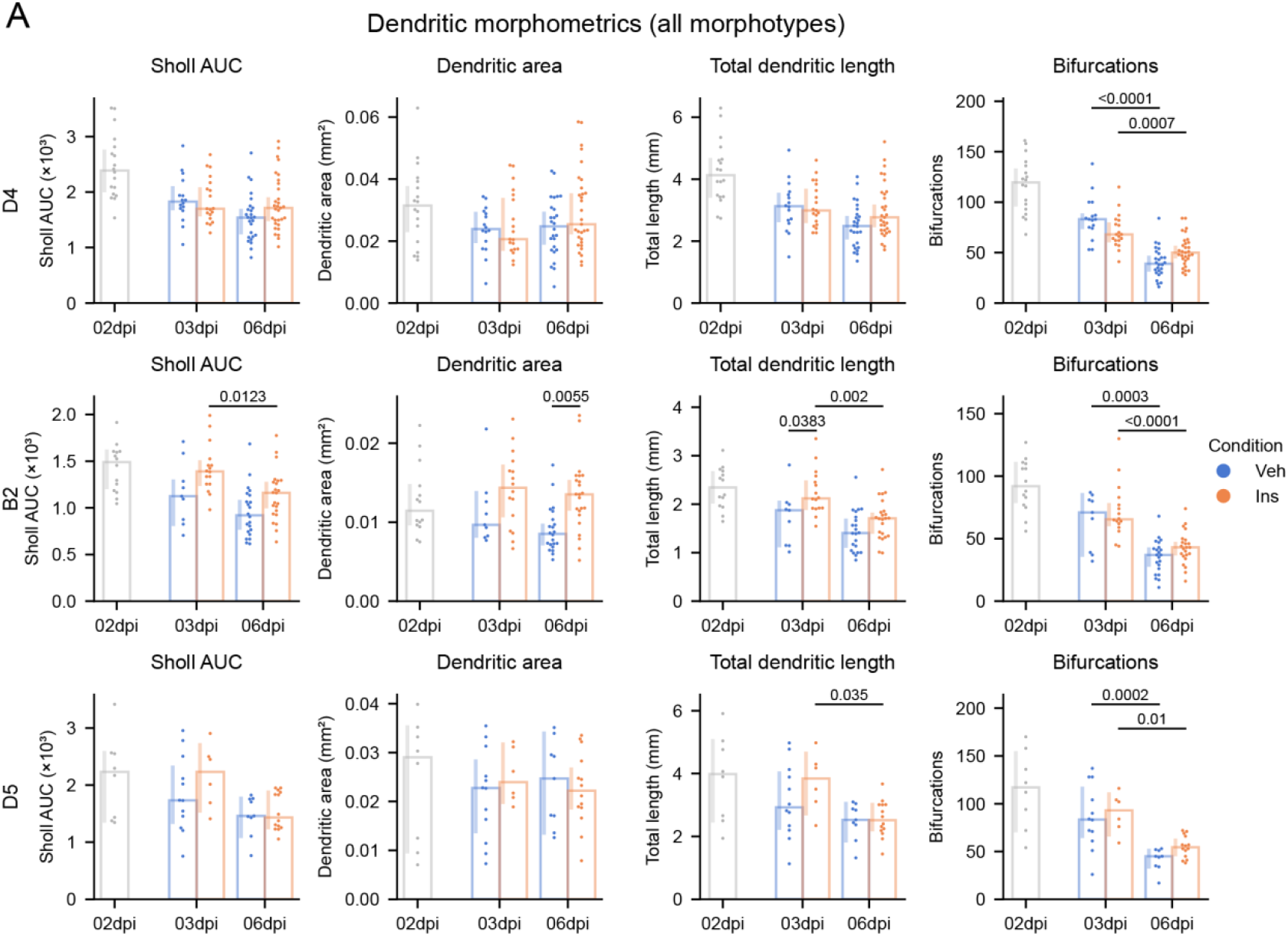
Dendritic morphometrics of insulin-treated gfi1ab+ RGC morphotypes. (A) Quantification of Sholl AUC, dendritic area, total dendritic length, and number of bifurcations for each morphotype at 2 dpi and following insulin or vehicle treatment at 3 and 6 dpi. Both vehicle- and insulin-treated cells exhibited dendritic pruning between 3dpi and 6dpi, as evidenced by the significant decrease in bifurcations. Overall trends point at an increased preservation of dendritic complexity with insulin-treatment compared with vehicles controls, particularly for the B2 morphotype. The 2 dpi data (grey) are taken as reference from Fig. 2 for comparison with the insulin- and vehicle-treated groups. Data presented as median ±95% confidence interval. Each dot represents one cell (N=40-47 cells from n=5-8 fish at 3 dpi, N=71-74 cells from n = 10 fish at 6dpi). One-way Kruskal-Wallis ANOVA with multiple comparisons followed by Mann–Whitney post hoc tests; p values are shown in the figure. Abbreviations: Area under the curve (AUC); days post-injury (dpi).

**Figure S5.**
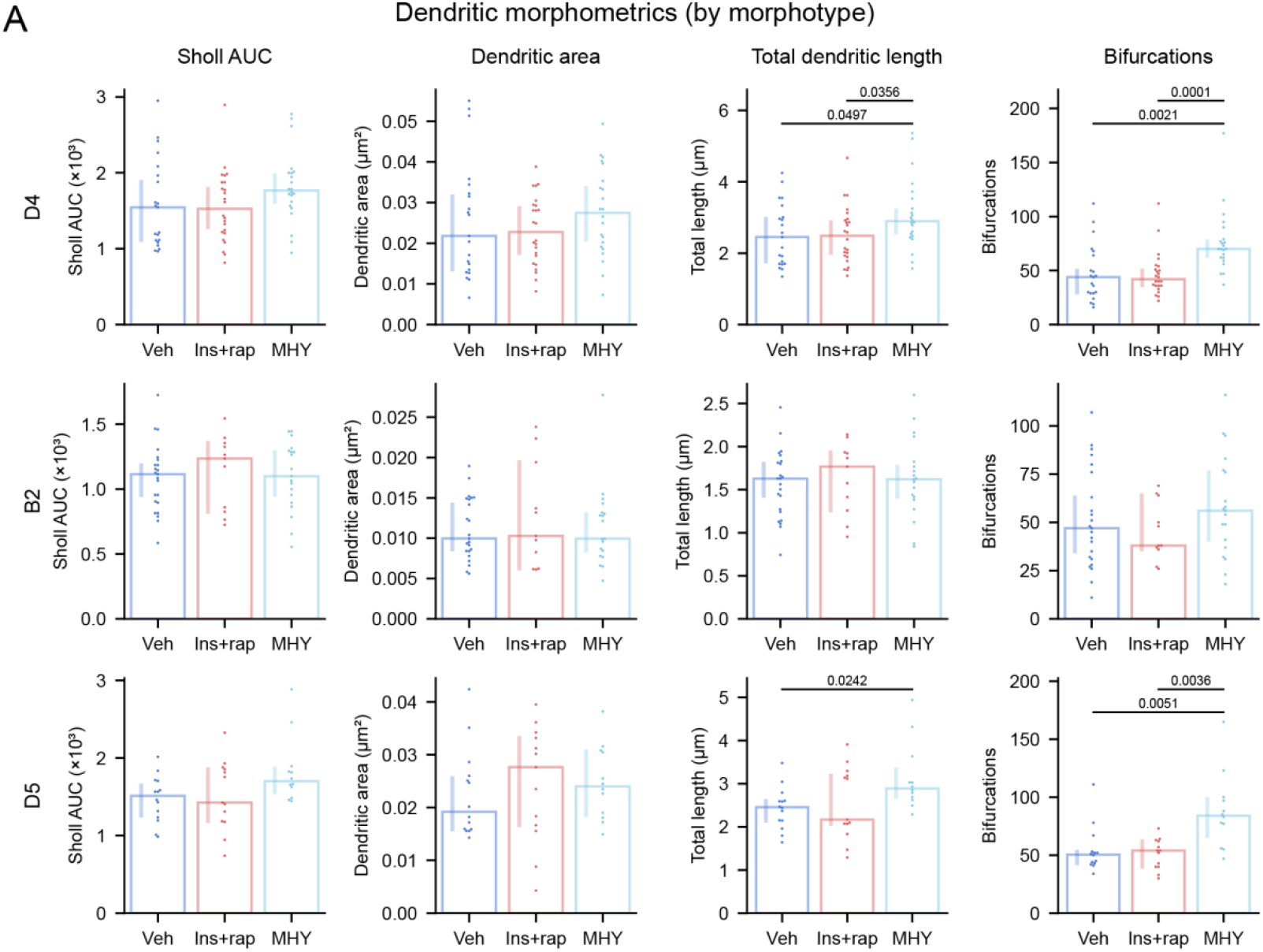
Dendritic morphometrics of insulin with rapamycin or MHY1485 treated gfi1ab+ RGC morphotypes. A) Quantification of dendritic morphometrics (Sholl AUC, dendritic area, total dendritic length, and number of bifurcations) at 6dpi following vehicle, combined insulin and rapamycin or MHY1485 treatment. Cells treated with insulin and rapamycin exhibit a dendritic complexity comparable to vehicle controls across all morphotypes. MHY1485-treated cells showed preservation of dendritic complexity relative to vehicle controls at 6 dpi, particularly in the number of bifurcations across all morphotypes. Data presented as median ±95% confidence interval. Each dot represents one cell (N=52-63 cells sampled from n = 7 fish per group). One-way Kruskal-Wallis ANOVA with multiple comparisons followed by Mann–Whitney post hoc tests; p values are shown in the figure. Abbreviations: Area under the curve (AUC); days post-injury (dpi); insulin (Ins); rapamycin (Rap); vehicle (Veh)

## Notes

### Competing Interest Statement

The authors have declared no competing interest.

## References

1. Williams, P.R., Benowitz, L.I., Goldberg, J.L., and He, Z. (2020). Axon Regeneration in the Mammalian Optic Nerve. Annu. Rev. Vis. Sci. 6, 195–213. 10.1146/annurev-vision-022720-094953.

2. Fawcett, J.W. (2020). The Struggle to Make CNS Axons Regenerate: Why Has It Been so Difficult? Neurochem. Res. 45, 144–158. 10.1007/s11064-019-02844-y.

3. Varadarajan, S.G., Hunyara, J.L., Hamilton, N.R., Kolodkin, A.L., and Huberman, A.D. (2022). Central nervous system regeneration. Cell 185, 77–94. 10.1016/j.cell.2021.10.029.

4. Takata, Y., Yamanaka, H., Nakagawa, H., and Takada, M. (2023). Morphological changes of large layer V pyramidal neurons in cortical motor-related areas after spinal cord injury in macaque monkeys. Scientific Reports 2023 13:1 13, 82-. 10.1038/s41598-022-26931-3.

5. Tseng, G.F., and Hu, M.E. (1996). Axotomy Induces Retraction of the Dendritic Arbor of Adult Rat Rubrospinal Neurons. Acta Anat. (Basel). 155, 184–193. 10.1159/000147803.

6. Leung, C.K.S., Weinreb, R.N., Li, Z.W., Liu, S., Lindsey, J.D., Choi, N., Liu, L., Cheung, C.Y. lui, Ye, C., Qiu, K., et al. (2011). Long-term in vivo imaging and measurement of dendritic shrinkage of retinal ganglion cells. Invest. Ophthalmol. Vis. Sci. 52, 1539–1547. 10.1167/iovs.10-6012.

7. Kalesnykas, G., Oglesby, E.N., Zack, D.J., Cone, F.E., Steinhart, M.R., Tian, J., Pease, M.E., and Quigley, H.A. (2012). Retinal Ganglion Cell Morphology after Optic Nerve Crush and Experimental Glaucoma. Invest. Ophthalmol. Vis. Sci. 53, 3847–3857. 10.1167/IOVS.12-9712.

8. Di Polo, A. (2015). Dendrite pathology and neurodegeneration: Focus on mTOR. Neural Regen. Res. 10, 559–561. 10.4103/1673-5374.155421.

9. Morquette, B., Morquette, P., Agostinone, J., Feinstein, E., McKinney, R.A., Kolta, A., and Di Polo, A. (2015). REDD2-mediated inhibition of mTOR promotes dendrite retraction induced by axonal injury. Cell Death Differ. 22, 612–625. 10.1038/cdd.2014.149.

10. Agostinone, J., Alarcon-Martinez, L., Gamlin, C., Yu, W.Q., Wong, R.O.L., and Di Polo, A. (2018). Insulin signalling promotes dendrite and synapse regeneration and restores circuit function after axonal injury. Brain 141, 1963–1980. 10.1093/brain/awy142.

11. El Hajji, S., Shiga, Y., Belforte, N., Solorio, Y.C., Tastet, O., D’Onofrio, P., Dotigny, F., Prat, A., Arbour, N., Fortune, B., et al. (2024). Insulin restores retinal ganglion cell functional connectivity and promotes visual recovery in glaucoma. Sci. Adv. 10, 5722. 10.1126/SCIADV.ADL5722.

12. Beckers, A., Van Dyck, A., Bollaerts, I., Van houcke, J., Lefevere, E., Andries, L., Agostinone, J., Van Hove, I., Di Polo, A., Lemmens, K., et al. (2019). An Antagonistic Axon-Dendrite Interplay Enables Efficient Neuronal Repair in the Adult Zebrafish Central Nervous System. Mol. Neurobiol. 56, 3175–3192. 10.1007/s12035-018-1292-5.

13. Chung, S.H., Awal, M.R., Shay, J., McLoed, M.M., Mazur, E., and Gabel, C. V. (2016). Novel DLK-independent neuronal regeneration in Caenorhabditis elegans shares links with activity-dependent ectopic outgrowth. Proc. Natl. Acad. Sci. U. S. A. 113, 2852–2860. 10.1073/pnas.1600564113.

14. Drummond, E.S., Rodger, J., Penrose, M., Robertson, D., Hu, Y., and Harvey, A.R. (2014). Effects of intravitreal injection of a Rho-GTPase inhibitor (BA-210), or CNTF combined with an analogue of cAMP, on the dendritic morphology of regenerating retinal ganglion cells. Restor. Neurol. Neurosci. 32, 391–402. 10.3233/RNN-130360.

15. Santos, J.R.F., Li, C., Andries, L., Masin, L., Nuttin, B., Reinhard, K., Moons, L., Cuntz, H., and Farrow, K. (2025). Developmental trajectories predict dendritic remodeling after injury. iScience 28, 113373. 10.1016/j.isci.2025.113373.

16. Putti, E., Faini, G., Thanh-Mai Dang, J., Savaliya, J.H., Eggeler, F., Duroure, K., Vougny, J., Ortiz-Álvarez, G., Pujades, C., Bahl, A., et al. (2026). Lrrn-mediated retinal ganglion cell targeting drives visual circuit assembly for brightness and contrast detection. Sci. Adv. 12, 4585. 10.1126/sciadv.adz4585.

17. Mangrum, W.I., Dowling, J.E., and Cohen, E.D. (2002). A morphological classification of ganglion cells in the zebrafish retina. Vis. Neurosci. 19, 767–779. 10.1017/S0952523802196076.

18. Robles, E., Laurell, E., and Baier, H. (2014). The Retinal Projectome Reveals Brain-Area-Specific Visual Representations Generated by Ganglion Cell Diversity. Current Biology 24, 2085–2096. 10.1016/j.cub.2014.07.080.

19. Van houcke, J., Geeraerts, E., Vanhunsel, S., Beckers, A., Noterdaeme, L., Christiaens, M., Bollaerts, I., De Groef, L., and Moons, L. (2019). Extensive growth is followed by neurodegenerative pathology in the continuously expanding adult zebrafish retina. Biogerontology 20, 109–125. 10.1007/s10522-018-9780-6.

20. Dhara, S.P., Rau, A., Flister, M.J., Recka, N.M., Laiosa, M.D., Auer, P.L., and Udvadia, A.J. (2019). Cellular reprogramming for successful CNS axon regeneration is driven by a temporally changing cast of transcription factors. Scientific Reports 2019 9:1 9, 1–12. 10.1038/s41598-019-50485-6.

21. Diekmann, H., Kalbhen, P., and Fischer, D. (2015). Active mechanistic target of rapamycin plays an ancillary rather than essential role in zebrafish CNS axon regeneration. Front. Cell. Neurosci. 9, 251. 10.3389/fncel.2015.00251.

22. Zhang, A., Bergmans, S., Dyck, A. Van, Beckers, A., Moons, L., and Masin, L. (2025). Successful axonal regeneration is associated with intraneuronal metabolic reprogramming. iScience 28, 113631. 10.1016/J.ISCI.2025.113631.

23. Lipton, J.O., and Sahin, M. (2014). The Neurology of mTOR. Neuron 84, 275–291. 10.1016/j.neuron.2014.09.034.

24. Rui, Y., Myers, K.R., Yu, K., Wise, A., De Blas, A.L., Hartzell, H.C., and Zheng, J.Q. (2013). Activity-dependent regulation of dendritic growth and maintenance by glycogen synthase kinase 3β. Nat. Commun. 4, 1–13. 10.1038/NCOMMS3628;TECHMETA.

25. Santina, L. Della, Inman, D.M., Lupien, C.B., Horner, P.J., and Wong, R.O.L. (2013). Differential progression of structural and functional alterations in distinct retinal ganglion cell types in a mouse model of glaucoma. Journal of Neuroscience 33, 17444–17457. 10.1523/JNEUROSCI.5461-12.2013.

26. Williams, P.A., Howell, G.R., Barbay, J.M., Braine, C.E., Sousa, G.L., John, S.W.M., and Morgan, J.E. (2013). Retinal Ganglion Cell Dendritic Atrophy in DBA/2J Glaucoma. PLoS One 8, e72282. 10.1371/JOURNAL.PONE.0072282.

27. Tribble, J.R., Vasalauskaite, A., Redmond, T., Young, R.D., Hassan, S., Fautsch, M.P., Sengpiel, F., Williams, P.A., and Morgan, J.E. (2019). Midget retinal ganglion cell dendritic and mitochondrial degeneration is an early feature of human glaucoma. Brain Commun. 1. 10.1093/BRAINCOMMS/FCZ035.

28. Claes, M., Santos, J.R.F., Masin, L., Cools, L., Davis, B.M., Arckens, L., Farrow, K., De Groef, L., and Moons, L. (2021). A fair assessment of evaluation tools for the murine microbead occlusion model of glaucoma. Int. J. Mol. Sci. 22, 5633. 10.3390/ijms22115633.

29. Zolessi, F.R., Poggi, L., Wilkinson, C.J., Chien, C. Bin, and Harris, W.A. (2006). Polarization and orientation of retinal ganglion cells in vivo. Neural Development 2006 1:1 1, 2-. 10.1186/1749-8104-1-2.

30. Wingate, R.J.T., and Thompson, I.D. (1995). The morphological development of mammalian retinal ganglion cells. Prog. Retin. Eye Res. 14, 413–435. 10.1016/1350-9462(94)00013-6.

31. Goldberg, J.L., Klassen, M.P., Hua, Y., and Barres, B.A. (2002). Amacrine-Signaled Loss of Intrinsic Axon Growth Ability by Retinal Ganglion Cells. Science (1979). 296, 1860–1864. 10.1126/science.1068428.

32. Berry, M., Ahmed, Z., Morgan-Warren, P., Fulton, D., and Logan, A. (2016). Prospects for mTOR-mediated functional repair after central nervous system trauma. Neurobiol. Dis. 85, 99–110. 10.1016/J.NBD.2015.10.002.

33. Duan, X., Qiao, M., Bei, F., Kim, I.J., He, Z., and Sanes, J.R. (2015). Subtype-Specific regeneration of retinal ganglion cells following axotomy: Effects of osteopontin and mtor signaling. Neuron 85, 1244–1256. 10.1016/j.neuron.2015.02.017.

34. Liu, K., Lu, Y., Lee, J.K., Samara, R., Willenberg, R., Sears-Kraxberger, I., Tedeschi, A., Park, K.K., Jin, D., Cai, B., et al. (2010). PTEN deletion enhances the regenerative ability of adult corticospinal neurons. Nature Neuroscience 2010 13:9 13, 1075–1081. 10.1038/nn.2603.

35. Nieuwenhuis, B., Barber, A.C., Evans, R.S., Pearson, C.S., Fuchs, J., MacQueen, A.R., Erp, S. van, Haenzi, B., Hulshof, L.A., Osborne, A., et al. (2020). PI 3-kinase delta enhances axonal PIP3 to support axon regeneration in the adult CNS. EMBO Mol. Med. 12, e11674. 10.15252/EMMM.201911674.

36. Park, K.K., Liu, K., Hu, Y., Smith, P.D., Wang, C., Cai, B., Xu, B., Connolly, L., Kramvis, I., Sahin, M., et al. (2008). Promoting axon regeneration in the adult CNS by modulation of the PTEN/mTOR pathway. Science (1979). 322, 963–966. 10.1126/science.1161566.

37. Choi, Y.J., Park, Y.J., Park, J.Y., Jeong, H.O., Kim, D.H., Ha, Y.M., Kim, J.M., Song, Y.M., Heo, H.S., Yu, B.P., et al. (2012). Inhibitory Effect of mTOR Activator MHY1485 on Autophagy: Suppression of Lysosomal Fusion. PLoS One 7, e43418. 10.1371/JOURNAL.PONE.0043418.

38. Sun, L., Morikawa, K., Sogo, Y., and Sugiura, Y. (2021). MHY1485 enhances X-irradiation-induced apoptosis and senescence in tumor cells. J. Radiat. Res. 62. 10.1093/jrr/rrab057.

39. Lu, F., Leach, L.L., and Gross, J.M. (2022). mTOR activity is essential for retinal pigment epithelium regeneration in zebrafish. PLoS Genet. 18. 10.1371/journal.pgen.1009628.

40. Luo, B., Cheng, T., Xiang, Y., Sun, S., Liu, Y., Yan, J., Zhang, Y., Cao, Y., Liu, X., and Pei, R. (2025). Promoting retinal ganglion cell regeneration with targeted liposome-based delivery of MHY1485 for optic nerve repair. Journal of Controlled Release 383. 10.1016/j.jconrel.2025.113778.

41. Cook, N.E., McGovern, M.R., Zaman, T., Lundin, P.M., and Vaughan, R.A. (2024). Effect of mTORC Agonism via MHY1485 with and without Rapamycin on C2C12 Myotube Metabolism. Int. J. Mol. Sci. 25. 10.3390/ijms25136819.

42. Lewis, T.L., Turi, G.F., Kwon, S.-K., Losonczy, A., and Polleux, F. (2016). Progressive Decrease of Mitochondrial Motility during Maturation of Cortical Axons In Vitro and In Vivo. Current Biology 26, 2602–2608. 10.1016/j.cub.2016.07.064.

43. Koseki, H., Donegá, M., Lam, B.Y.H., Petrova, V., van Erp, S., Yeo, G.S.H., Kwok, J.C.F., Ffrench-Constant, C., Eva, R., and Fawcett, J.W. (2017). Selective rab11 transport and the intrinsic regenerative ability of CNS axons. Elife 6. 10.7554/ELIFE.26956.

44. Eva, R., Koseki, H., Kanamarlapudi, V., and Fawcett, J.W. (2017). EFA6 regulates selective polarised transport and axon regeneration from the axon initial segment. J. Cell Sci. 130, 3663–3675. 10.1242/JCS.207423/VIDEO-8.

45. Lee, C.C., Huang, C.C., Wu, M.Y., and Hsu, K. Sen (2005). Insulin Stimulates Postsynaptic Density-95 Protein Translation via the Phosphoinositide 3-Kinase-Akt-Mammalian Target of Rapamycin Signaling Pathway. Journal of Biological Chemistry 280, 18543–18550. 10.1074/JBC.M414112200.

46. Chiu, S.L., and Cline, H.T. (2010). Insulin receptor signaling in the development of neuronal structure and function. Neural Dev. 5, 7-. 10.1186/1749-8104-5-7/FIGURES/4.

47. Gralle, M. (2017). The neuronal insulin receptor in its environment. J. Neurochem. 140, 359–367. 10.1111/JNC.13909;PAGE:STRING:ARTICLE/CHAPTER.

48. Costa-Mattioli, M., Sossin, W.S., Klann, E., and Sonenberg, N. (2009). Translational Control of Long-Lasting Synaptic Plasticity and Memory. Neuron 61, 10–26. 10.1016/J.NEURON.2008.10.055.

49. Dupraz, S., Grassi, D., Karnas, D., Nieto Guil, A.F., Hicks, D., and Quiroga, S. (2013). The Insulin-Like Growth Factor 1 Receptor Is Essential for Axonal Regeneration in Adult Central Nervous System Neurons. PLoS One 8. 10.1371/journal.pone.0054462.

50. Liu, Y., Wang, X., Li, W., Zhang, Q., Li, Y., Zhang, Z., Zhu, J., Chen, B., Williams, P.R., Zhang, Y., et al. (2017). A Sensitized IGF1 Treatment Restores Corticospinal Axon-Dependent Functions. Neuron 95. 10.1016/j.neuron.2017.07.037.

51. Terenzio, M., Koley, S., Samra, N., Rishal, I., Zhao, Q., Sahoo, P.K., Urisman, A., Marvaldi, L., Oses-Prieto, J.A., Forester, C., et al. (2018). Locally translated mTOR controls axonal local translation in nerve injury. Science (1979). 359, 1416–1421. 10.1126/SCIENCE.AAN1053/SUPPL_FILE/AAN1053_TERENZIO_SM_TABLE_S2.XLSX.

52. Poulopoulos, A., Murphy, A.J., Ozkan, A., Davis, P., Hatch, J., Kirchner, R., and Macklis, J.D. (2019). Subcellular transcriptomes and proteomes of developing axon projections in the cerebral cortex. Nature 565, 356–360. 10.1038/s41586-018-0847-y.

53. Altas, B., Romanowski, A.J., Bunce, G.W., and Poulopoulos, A. (2022). Neuronal mTOR Outposts: Implications for Translation, Signaling, and Plasticity. Front. Cell. Neurosci. 0, 134. 10.3389/FNCEL.2022.853634.

54. Lim, J.H.A., Stafford, B.K., Nguyen, P.L., Lien, B. V., Wang, C., Zukor, K., He, Z., and Huberman, A.D. (2016). Neural activity promotes long-distance, target-specific regeneration of adult retinal axons. Nat. Neurosci. 19, 1073–1084. 10.1038/nn.4340.

55. Beckers, A., Bergmans, S., Van Dyck, A., and Moons, L. (2023). Analysis of Axonal Regrowth and Dendritic Remodeling After Optic Nerve Crush in Adult Zebrafish. In Methods in Molecular Biology 10.1007/978-1-0716-3012-9_9.

56. Kölsch, Y., Hahn, J., Sappington, A., Stemmer, M., Fernandes, A.M., Helmbrecht, T.O., Lele, S., Butrus, S., Laurell, E., Arnold-Ammer, I., et al. (2021). Molecular classification of zebrafish retinal ganglion cells links genes to cell types to behavior. Neuron 109, 645-662.e9. 10.1016/j.neuron.2020.12.003.

57. Kölsch, Y., Hahn, J., Sappington, A., Stemmer, M., Fernandes, A.M., Helmbrecht, T.O., Lele, S., Butrus, S., Laurell, E., Arnold-Ammer, I., et al. (2021). Molecular classification of zebrafish retinal ganglion cells links genes to cell types to behavior. Neuron 109, 645-662.e9. 10.1016/J.NEURON.2020.12.003.

58. Beckers, A., Van Dyck, A., Bollaerts, I., Van houcke, J., Lefevere, E., Andries, L., Agostinone, J., Van Hove, I., Di Polo, A., Lemmens, K., et al. (2019). An Antagonistic Axon-Dendrite Interplay Enables Efficient Neuronal Repair in the Adult Zebrafish Central Nervous System. Mol. Neurobiol. 56, 3175–3192. 10.1007/S12035-018-1292-5/FIGURES/8.

59. Fang, Z., Liu, X., and Peltz, G. (2023). GSEApy: a comprehensive package for performing gene set enrichment analysis in Python. Bioinformatics 39. 10.1093/BIOINFORMATICS/BTAC757.

60. Arshadi, C., Günther, U., Eddison, M., Harrington, K.I.S., and Ferreira, T.A. (2021). SNT: a unifying toolbox for quantification of neuronal anatomy. Nat. Methods 18. 10.1038/s41592-021-01105-7.

61. Arnaudon, A., Berchet, A., Courcol, J.-D., Coste, B., Gevaert, M., Kanari, L., Sanin, A., Palacios, J., Vanherpe, L., and Zisis, E. NeuroM. 10.5281/ZENODO.10630119.

62. Sümbül, U., Song, S., McCulloch, K., Becker, M., Lin, B., Sanes, J.R., Masland, R.H., and Seung, H.S. (2014). A genetic and computational approach to structurally classify neuronal types. Nat. Commun. 5, 1–12. 10.1038/NCOMMS4512;TECHMETA.

