## Supplementary_figures_and_legends for "Repressed mTORC1 signaling and transient dendritic pruning support axonal regeneration"

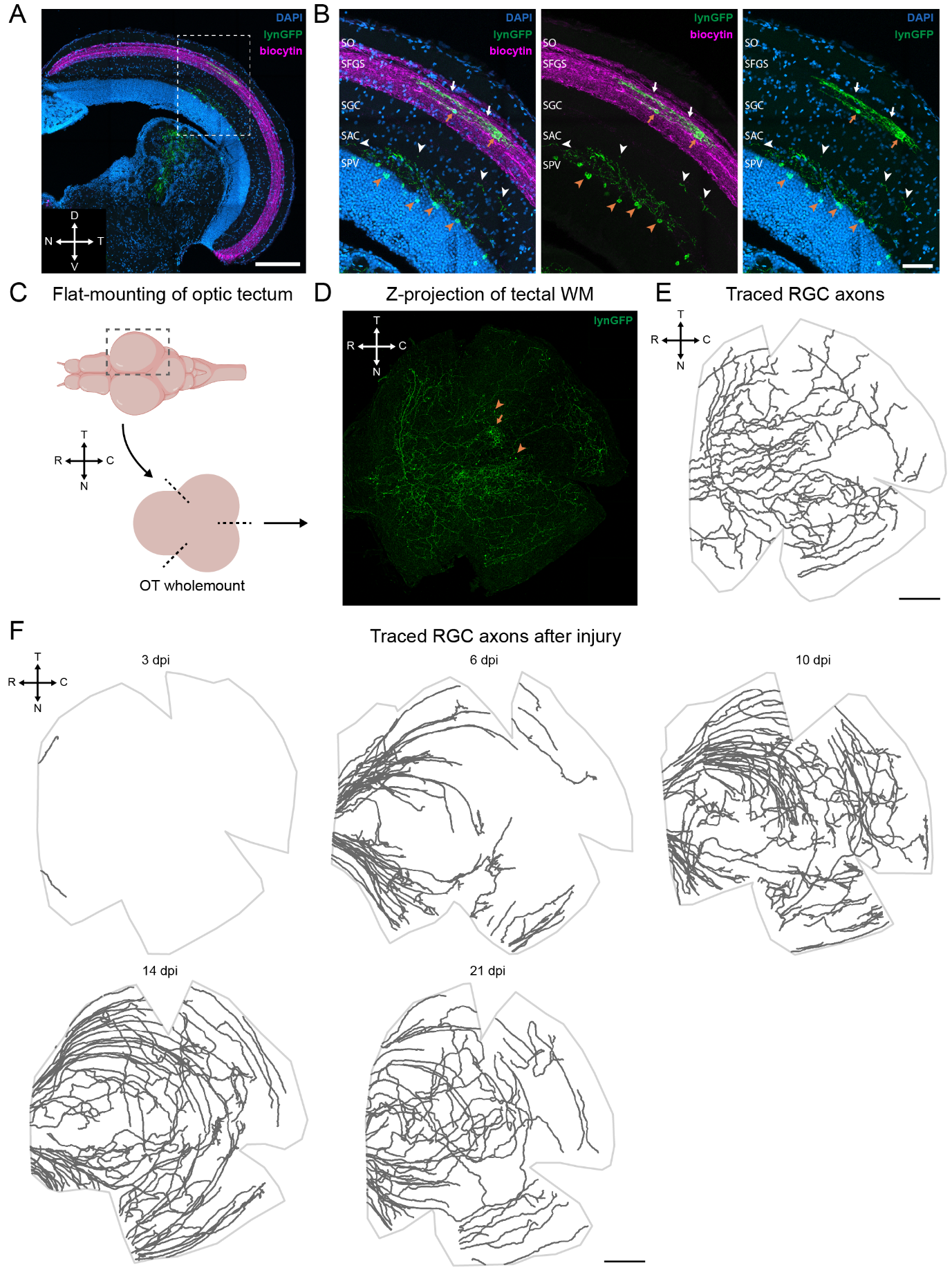

#### Figure S1. Tectal innervation pattern of gfi1ab+ RGCs and reinnervation timeline after injury

(A) Representative coronal brain cryosections from anterogradely traced fish labelled for DAPI, lynGFP and biocytin showing innervation of all RGCs (biocytin) and *gfi1ab*+ RGC axons in the dorsomedial optic tectum (white box) under non-injured conditions. Scale bar = 200 µm.

(B) Magnified views of the coronal brain section reveal localized innervation by *gfi1ab*+ axons primarily in the SFGS (white arrows). Tectal neurons in the SFGS near the axon terminals are also sparsely labelled (orange arrows), possibly indicating synapses with the incoming RGC axons. Sparser innervation, mostly running perpendicular to the sectioning plane, is detectable also in the SGC and in the SAC (white arrowheads). Here, RGC axons appear in proximity with dendrites of labelled tectal neurons located in the SPV (orange arrowheads). See also panel D. Scale bar = 50 µm.

(C) Schematic representation of optic tectum whole-mount preparation. The right optic tectum was peeled from the brain, followed by three cuts at the nasal–ventral, nasal–dorsal, and temporal positions.

(D) Z-projection of a representative optic tectum wholemount showing innervation of *gfi1ab+* RGC axons to the medial optic tectum at non injured condition. Here, a small number of tectal neuron residing in the SGS are also labelled (orange arrow, see panel B). More abundant are neurons residing deeper in the SPV (orange arrowheads, see panel B). Scale bar = 200 µm.

(E) Manually traced *gfi1ab+* RGC axons showing innervation in the central tectum at non injured condition. Scale bar = 200 µm.

F) Representative reconstructions of manually traced *gfi1ab+* RGC axons showing the progressive reinnervation of the tectum after optic nerve injury. At 3 dpi, only rare pioneering *gfi1ab+* RGC axons are visible the optic tectum. By 6 dpi, a subset of labeled axons enters the optic tectum from the nasal and temporal sides and navigates toward targets in the central tectum. At 10 dpi, the majority of labeled axons navigate toward their target region. By 14 dpi, axons have reached the target in the central tecum; however, further refinement of innervation could take place after this timepoint. Scale bar = 200 µm.

Abbreviations: caudal (C); choline acetyltransferase (ChAT); dorsal (D); days post-injury (dpi); inner plexiform layer (IPL); nasal (N); OT (optic tectum); retinal ganglion cell (RGC); rostral (R); temporal (T); ventral (V); whole-mount (WM).

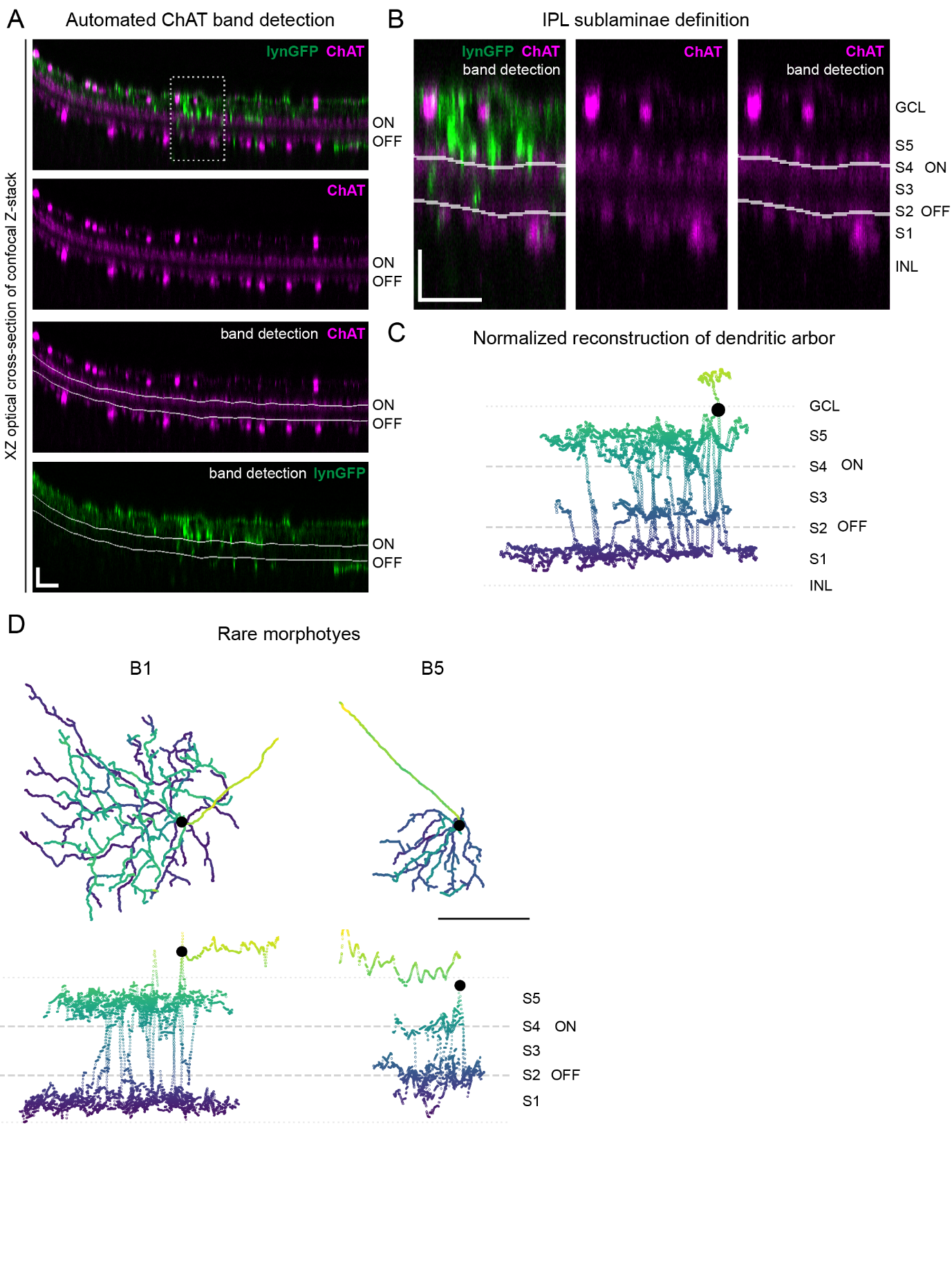

#### Figure S2. Normalization of RGC dendrite depth and representation of rare morphotypes

A) Optical section of a representative confocal stack used to image gfi1ab+ cells after immunostaining for lynGFP (green) and choline acetyltransferase (ChAT, magenta). An automated deep-learning-based model is run to detect the position of ON and OFF ChAT bands (white annotations) throughout the stack to be used as reference to normalize the coordinates of dendritic RGCs. This step is critical to cancel-out distortion artefacts due to irregular retinal whole-mounting. See Methods for a deeper methodological detail. Scale bars (vertical and horizontal) = 20 µm.

B) Zoomed view of the optical section depicted in A. Chat band annotation are used to define IPL sublaminae, named S5 to S1 spanning from the GCL to the INL. S2 and S4 are defined in correspondence of the OFF and ON ChAT bands, respectively. Scale bars (vertical and horizontal) = 20 µm.

C) After registration of coordinates relative to the reference ChAT band, normalized dendrite reconstructions are plotted. These faithfully represent the lamination of RGC dendrites within the IPL and are used to classify the 3 main morphotypes, namely D4, B2 and D5.

D) En-face (XY, top) and orthogonal (XZ, bottom) dendritic reconstructions of representative rare B1 and B5 morphotypes. Although rarity prevents a quantitative characterization of their dendritic properties, B1 cells present large dendritic arbors stratified on S1 and S5, while B5 neurons present smaller arbors stratifying on S2 and S4. Scale bar = 100 µm.

Abbreviations: choline acetyltransferase (ChAT); ganglion cell layer (GCL); inner plexiform layer (IPL); inner nuclear layer (INL).

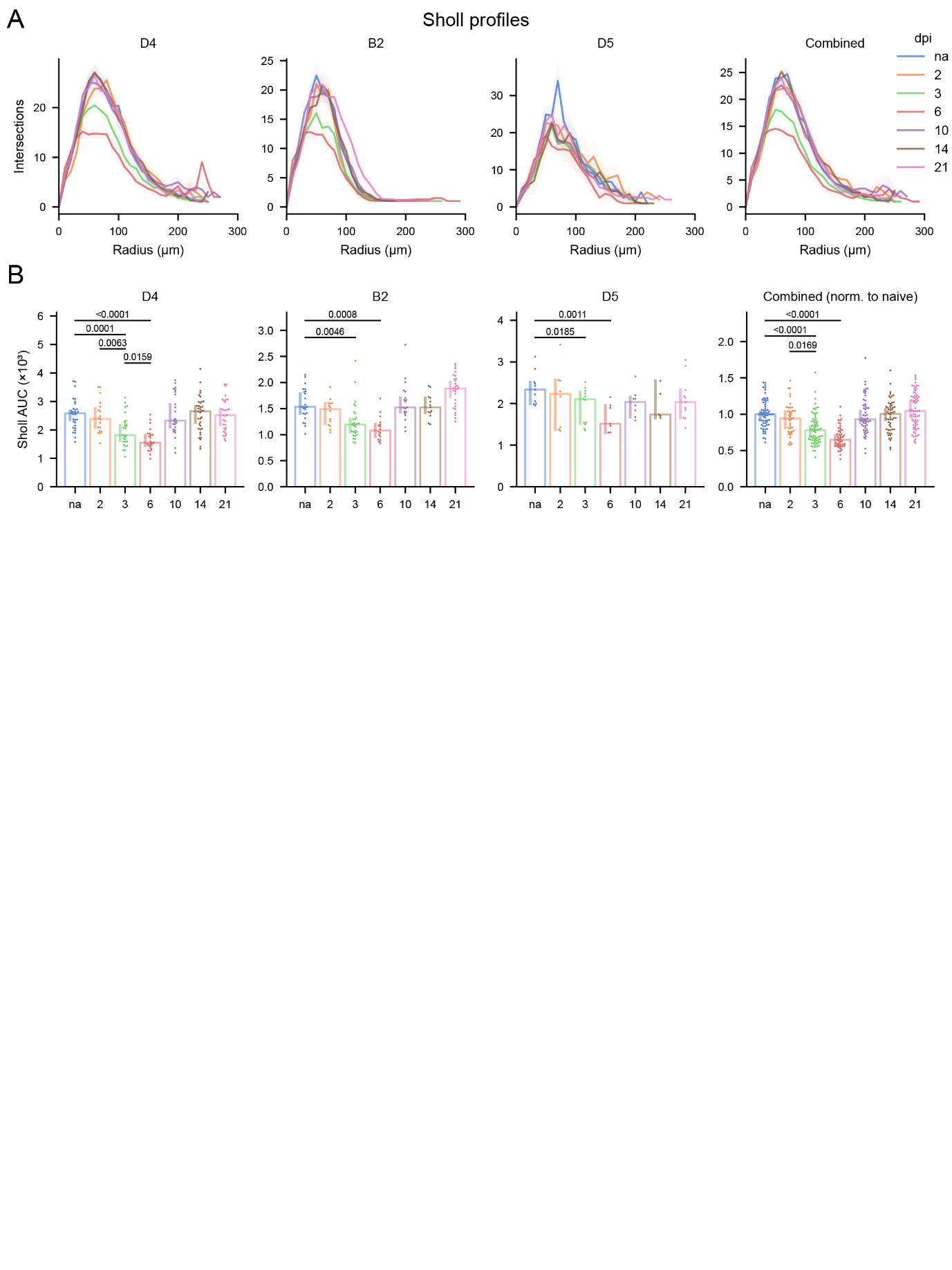

#### Figure S3. Sholl profiles of gfi1ab+ RGC morphotypes across the regeneration timeline.

(A) Representative Sholl profiles of D4, B2, and D5 morphotypes, as well as the combined cell population, across the regeneration timeline. Following injury, all morphotypes exhibit a transient reduction in dendritic complexity, evident from day 3 (green line) and most pronounced at day 6 (red line).
(B) Quantification of the area under the curve (AUC) of the Sholl profiles reveals significant dendritic pruning of RGCs at 3 and 6 dpi. Dendritic complexity is gradually restored by 10 dpi and further recovered through 21 dpi. Values are shown for all morphotypes individually as well as combined, after normalization to the mean of the corresponding uninjured morphotype

Data presented as median ±95% confidence interval. Each dot represents a cell (N=40-70 cells sampled across n=10 fish per timepoint). One-way Kruskal-Wallis ANOVA with multiple comparisons followed by Mann–Whitney post hoc tests; p values are shown in the figure.

Abbreviations: Area under the curve (AUC); days post-injury (dpi).

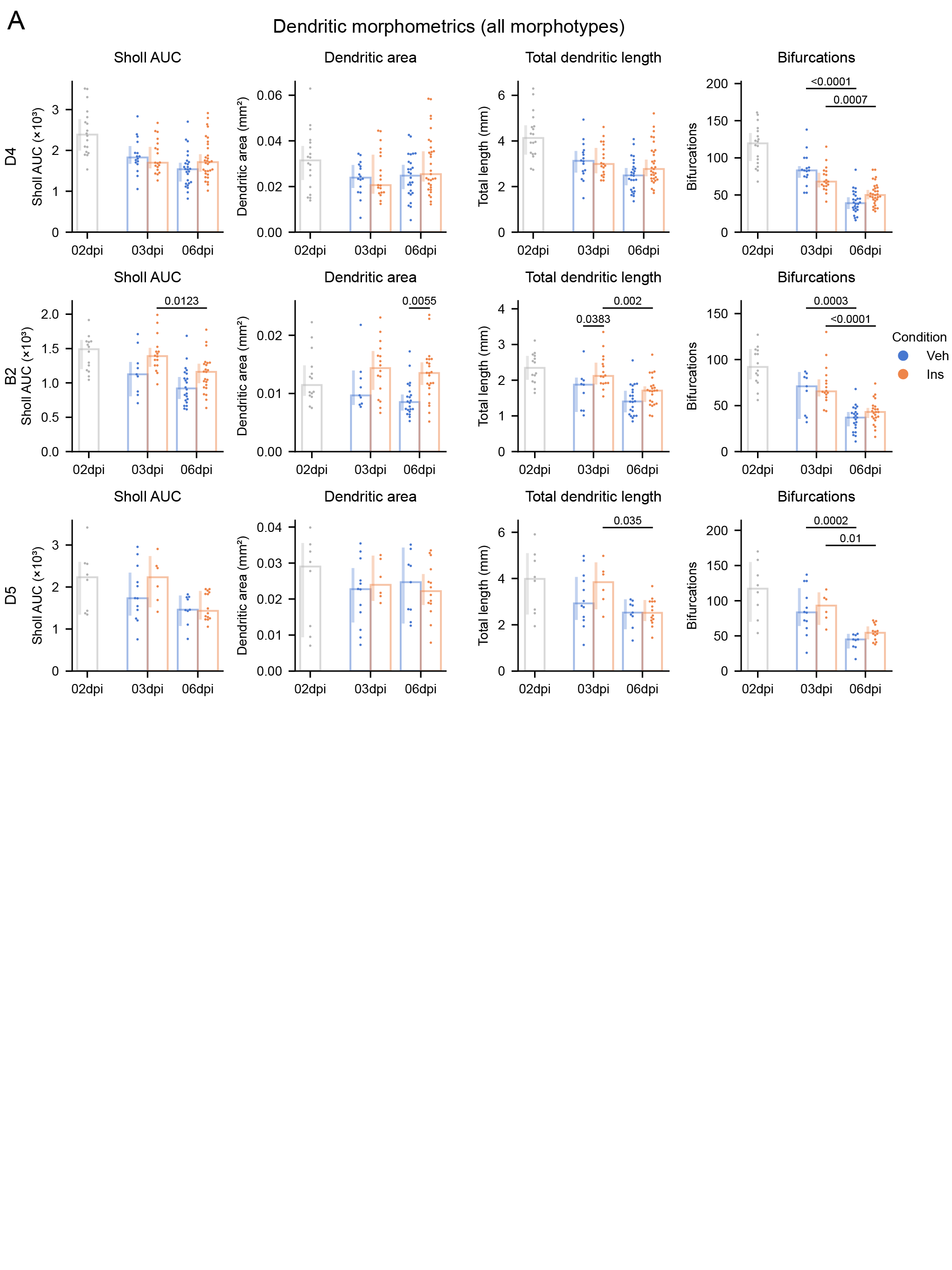

#### Figure S4. Dendritic morphometrics of insulin-treated gfi1ab+ RGC morphotypes

(A) Quantification of Sholl AUC, dendritic area, total dendritic length, and number of bifurcations for each morphotype at 2 dpi and following insulin or vehicle treatment at 3 and 6 dpi. Both vehicle- and insulin-treated cells exhibited dendritic pruning between 3dpi and 6dpi, as evidenced by the significant decrease in bifurcations. Overall trends point at an increased preservation of dendritic complexity with insulin-treatment compared with vehicles controls, particularly for the B2 morphotype. The 2 dpi data (grey) are taken as reference from Fig. 2 for comparison with the insulin- and vehicle-treated groups.

Data presented as median ±95% confidence interval. Each dot represents one cell (N=40-47 cells from n=5-8 fish at 3 dpi, N=71-74 cells from n = 10 fish at 6dpi). One-way Kruskal-Wallis ANOVA with multiple comparisons followed by Mann–Whitney post hoc tests; p values are shown in the figure.

Abbreviations: Area under the curve (AUC); days post-injury (dpi).

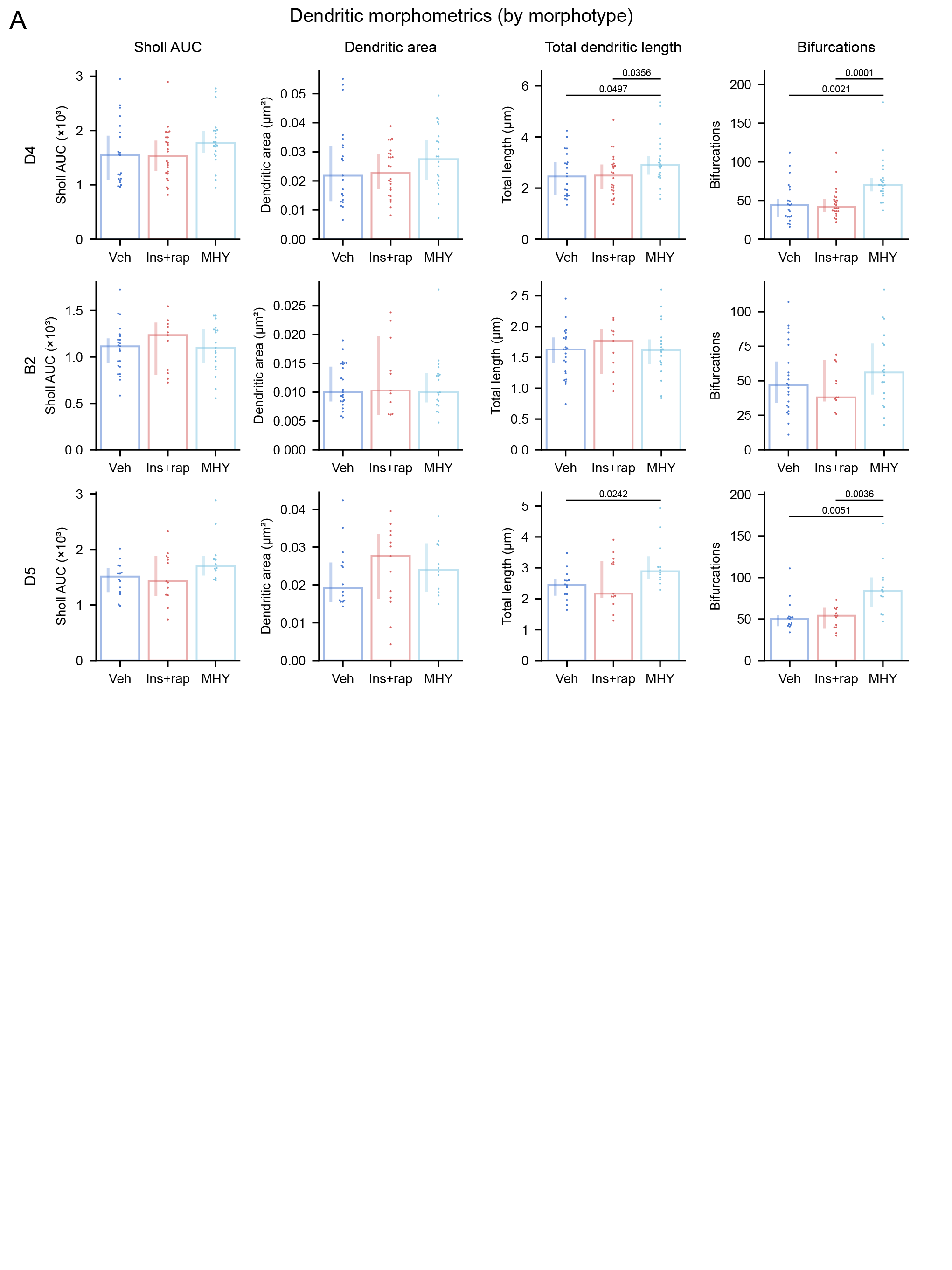

#### Figure S5. Dendritic morphometrics of insulin with rapamycin or MHY1485 treated gfi1ab+ RGC morphotypes

A) Quantification of dendritic morphometrics (Sholl AUC, dendritic area, total dendritic length, and number of bifurcations) at 6dpi following vehicle, combined insulin and rapamycin or MHY1485 treatment. Cells treated with insulin and rapamycin exhibit a dendritic complexity comparable to vehicle controls across all morphotypes. MHY1485-treated cells showed preservation of dendritic complexity relative to vehicle controls at 6 dpi, particularly in the number of bifurcations across all morphotypes.

Data presented as median ±95% confidence interval. Each dot represents one cell (N=52-63 cells sampled from n = 7 fish per group). One-way Kruskal-Wallis ANOVA with multiple comparisons followed by Mann–Whitney post hoc tests; p values are shown in the figure.

Abbreviations: Area under the curve (AUC); days post-injury (dpi); insulin (Ins); rapamycin (Rap); vehicle (Veh)
